# Single cell analysis of the cellular heterogeneity and interactions in the injured mouse spinal cord

**DOI:** 10.1101/2020.05.13.094854

**Authors:** Lindsay M Milich, James Choi, Christine Ryan, Stephanie L Yahn, Pantelis Tsoulfas, Jae K Lee

**Author notes:** These authors contributed equally (Lindsay M Milich, James Choi). Corresponding author: Jae K. Lee, PhD, University of Miami School of Medicine, Miami Project to Cure Paralysis, Department of Neurological Surgery, 1095 NW 14^th^ Terrace, LPLC 4-19, Miami, FL, 33136.

## Abstract

The wound healing process that occurs after spinal cord injury is critical for maintaining tissue homeostasis and limiting tissue damage, but eventually results in a scar-like environment that is not conducive to regeneration and repair. A better understanding of this dichotomy is critical to developing effective therapeutics that target the appropriate pathobiology, but a major challenge has been the large cellular heterogeneity that results in immensely complex cellular interactions. In this study, we used single cell RNA sequencing to assess virtually all cell types that comprise the mouse spinal cord injury site. In addition to discovering novel subpopulations, we used expression values of receptor-ligand pairs to identify signaling pathways that potentially drive specific cellular interactions during angiogenesis, gliosis, and fibrosis. Our dataset is a valuable resource that provides novel mechanistic insight into the pathobiology of not only spinal cord injury, but also other traumatic disorders of the CNS.

---

In the adult mammalian nervous system, a diverse population of glial and vascular cells are essential for optimal neuronal function. After spinal cord injury (SCI), this cellular diversity becomes even more heterogeneous by infiltrating leukocytes, which trigger a cascade of events collectively termed the wound healing process. The major research focus of the wound healing process after SCI has been its end-product, namely the formation of scar-like tissue in and around the lesion epicenter. However, we have a limited knowledge of the heterogeneity of the cells that comprise the injury site and how these cells interact during the wound healing process after SCI.

SCI triggers multiple processes that occur in a temporally defined manner. At 1 day post-injury (dpi), an innate immune response initiated by microglia is expanded by peripheral myeloid cells, mainly neutrophils and monocytes, which migrate to the injury site^1^. By 3dpi, most glial cells including astrocytes, microglia, and oligodendrocyte progenitor cells (OPCs), are at the peak of their proliferative state, resulting in recovery of cell number and initiating gliosis^2^. Concurrently, monocytes differentiate into macrophages that phagocytose cellular debris, acquire different phenotypic states, and activate perivascular fibroblasts to initiate fibrosis^3^. By 7dpi, the number of macrophages and fibroblasts have reached their peak, and the fibrotic scar begins to be surrounded by the astroglial scar, which limit their infiltration into the spinal cord parenchyma^4^. In addition, during these first 7 days, hypoxic conditions at the injury site promote angiogenesis and revascularization^5^. By 14dpi, gliosis, fibrosis, and angiogenesis have reached a more stable state^5–7^. Thus, the first 7 days after injury is highly dynamic, and provide a window of therapeutic intervention for multiple pathological processes, but we currently have a limited understanding of the cellular interactions that occur during this critical period.

One way to investigate cellular responses to injury is through transcriptomic profiling of cell types that comprise the injury site. Traditional methods of using FACS, or more recent methods utilizing immunoprecipitation of cell type-specific polyribosomes or nuclei have provided important insight into CNS pathobiology^8, 9^. However, these studies have been limited by the focus on a single cell type as well as the lack of information on subpopulation heterogeneity and different cell states. Recent advances in single cell RNA-sequencing (sc-RNAseq) technologies have circumvented these limitations by providing a more accurate depiction of the heterogeneous subpopulations that comprise a single cell type. In addition, many different cell types can be obtained and analyzed from a single tissue sample simultaneously, enabling examination of relationships and signaling mechanisms between multiple cell types.

In this study, we used sc-RNAseq to generate a single cell transcriptomics dataset of virtually all cell types that comprise the uninjured and injured spinal cord at 1, 3, and 7dpi. From this dataset, we were able to obtain unique molecular signatures of multiple cell types as well as their subpopulations present throughout the acute injury phase. By assessing expression of ligand-receptor pairs on different cell types, we were able to gain insight into potential signaling relationships that mediat angiogenesis, gliosis, and fibrosis. As the first sc-RNAseq analysis of the spinal cord injury site, our transcriptomic dataset will provide the field with novel and comprehensive insight into early spinal cord injury pathology as well as other traumatic injuries to the CNS.

## Results

### Molecular identification of immune, vascular, and glial cells in the acute injury site

To assess the cellular heterogeneity among all cell populations at the injury site, we obtained a total of 51,843 cells from uninjured and 1, 3, and 7dpi tissue, which resulted in a total of 15 distinct clusters when visualized on a UMAP plot (Fig. 1a, b). These 15 clusters represented all major cell types that are known to comprise the SCI site including microglia, monocytes, macrophages, neutrophils, dendritic cells, astrocytes, oligodendrocytes, OPCs, neurons, fibroblasts, pericytes, ependymal cells, and endothelial cells. These cell types were grouped into three categories, namely myeloid, vascular, and macroglia, for further analysis as described below. Neurons were excluded from the analysis because they were not expected to survive our dissociation protocol and thus subject to a selection bias. Lymphocytes were also excluded because they are primarily involved in autoimmunity after SCI^10^, which is beyond the scope of this study. Cell types pertaining to each cluster were identified using annotated lineage markers (Extended Data Fig. 1).

**Fig. 1.**
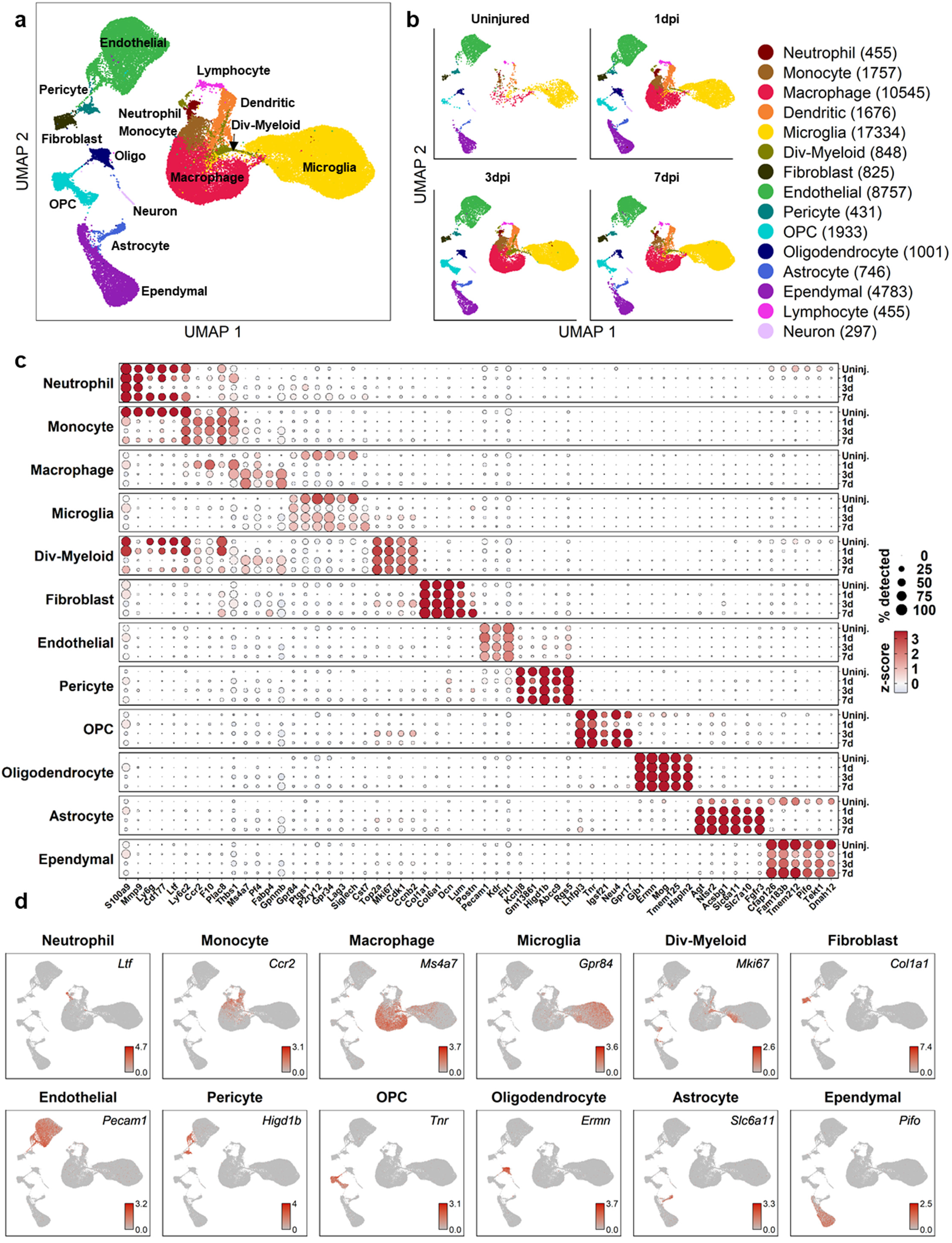
Transcriptomic identification of major cell types that comprise the spinal cord injury site at acute time points. (a) UMAP visualization of all cells from the uninjured, 1, 3, and 7dpi spinal cord displays all major cell types that are known to comprise the injury site. Clusters were identified based on expression of annotated genes for each cell type (Extended Data Fig. 1). (b) UMAP of each time point shows temporal progression of each major cell type. (c) Dot plot of highest differentially expressed genes (DEG) for each major cell type at each time point (right axis). Dot color intensity represents the z-score of expression values, and dot size represents the percentage of cells in each cell type expressing the DEG. (d) Expression pattern of the DEG that best identifies each cell type. n=2 biological replicates at each time point.

The highest differentially expressed genes (DEGs) provided a unique molecular signature for each cell type (Fig. 1c, d), which in most cases were different from canonical markers used in the literature. For example, the highest DEGs in OPCs were non-canonical genes such as *tnr* (Tenascin-R) and *lhfpl3* (lipoma HMGIG fusion partner), which displayed better specificity than canonical OPC markers such as *pdgfra* and *cspg4* that were expressed in multiple cell types (Extended Data Fig. 1). Interestingly, while certain marker genes were expressed both before and after injury, others such as *postn* in fibroblasts, changed expression in an injury-dependent manner. While *postn* was not expressed in uninjured fibroblasts, there was a graded increase as injury progressed (Fig. 1c). We validated this by genetic lineage tracing in Postn^EYFP^ mice, which showed EYFP cells present in the fibrotic scar and overlying meninges but absent in surrounding spinal cord tissue. (Extended Data Fig. 2). Taken together, our analysis of DEGs between major cell types uncovers highly specific molecular identifiers, many of which are non-canonical and display temporal specificity.

### Myeloid analysis reveals temporal changes in macrophage and microglial subtypes

To determine the heterogeneity within the myeloid population, clustering analysis was performed on myeloid cells and visualized on a separate UMAP (Fig. 2a, b), which revealed two large clusters corresponding to microglia and peripherally-derived myeloid cells as identified by annotated markers (Extended Data Fig. 3). We identified 6 microglial subtypes. Homeostatic microglia were identified based on its expression of several annotated markers of steady-state microglia, such as *p2ry12, siglech*, and *tmem119* (Fig. 2). As expected, homeostatic microglia were the predominant myeloid population in the uninjured spinal cord, but by 1dpi they were replaced by microglia subtypes similar to previously reported Disease-Associated Microglia (DAM) (Fig. 2b, c)^11^. We identified four DAM subtypes, which were labeled DAM-A to D. DAM-A was identified by high expression of the low density lipoprotein receptor *msr1* and low expression of the purinergic receptor *p2ry12*, and comprised 100% of microglia present at 1dpi. DAM-B and C expressed moderate level of *p2ry12* and low level of *msr1*, and their expression profile was similar to homeostatic microglia, suggesting an intermediate state as some DAM-A revert back to a more homeostatic state. DAM-D had low levels of both *msr1* and *p2ry12*, and was best identified by high expression of the growth factor *igf1* and the cholesterol-binding protein *apoe*. The last microglia subtype was identified as dividing microglia due to their high expression of cell cycle-related genes (*mki67* and *top2a*). Dividing microglia shared several DEGs with DAM-D such as *igf1, spp1*, and *fabp5* (Fig. 2e), and the fact that expansion of DAM-D from 3 to 7 dpi coincides with reduction in dividing microglia (Fig. 2c) suggests that dividing microglia may give rise to DAM-D after injury. A putative model depicting the relationship between DAM subtypes is illustrated in Extended Data Fig. 4a.

**Fig. 2.**
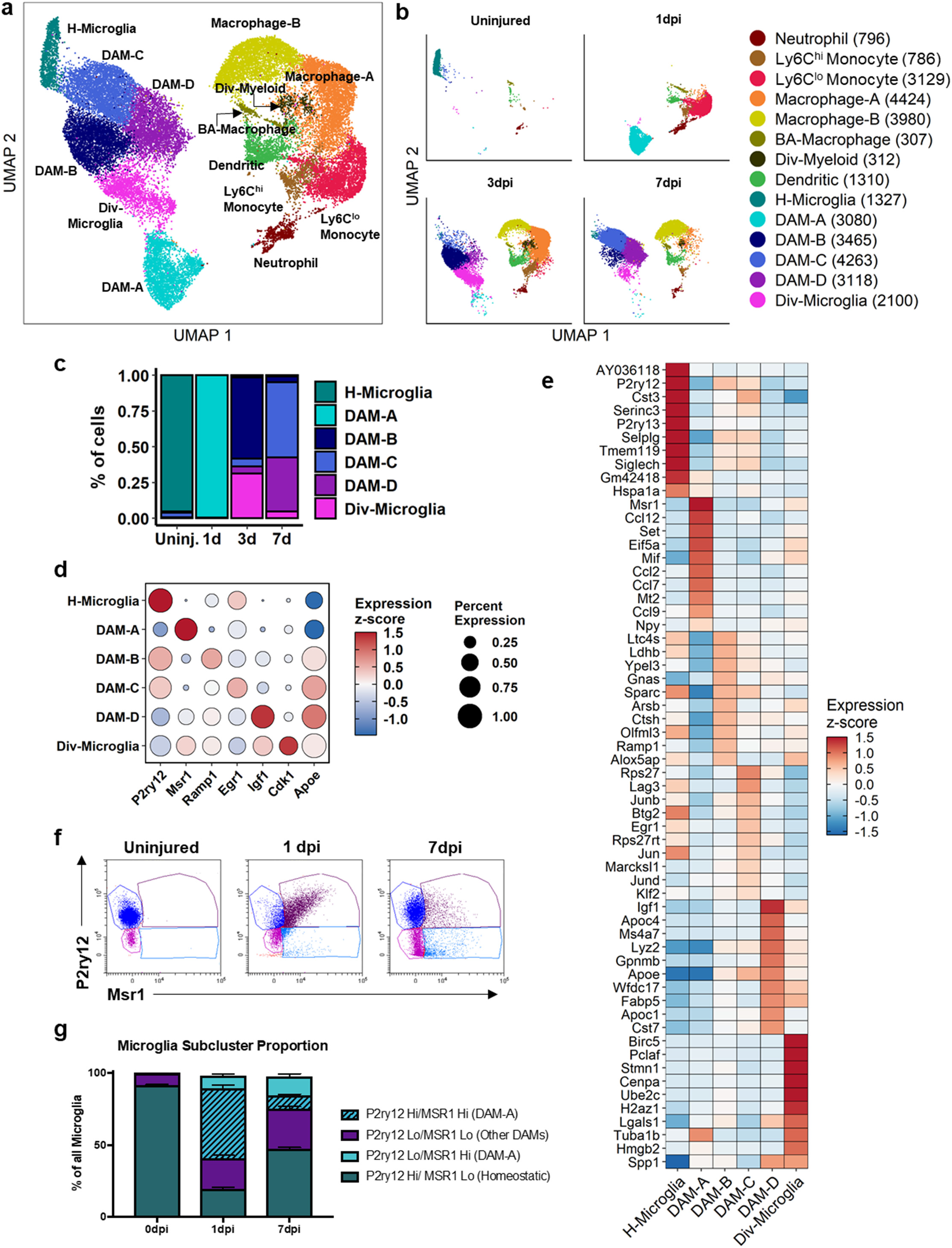
Molecular and temporal profile of microglia subtype heterogeneity acutely after SCI. (a) UMAP visualization of all myeloid cell clusters combined from uninjured, 1, 3, and 7dpi. Clusters were identified based on previously annotated cell types (Extended Data Fig. 3). Four subtypes of Disease-Associated Microglia (DAM) were labeled DAM-A to D. (b) UMAP of each time point displays temporal progression of each myeloid population. (c) Percent of each microglia subtype at each time point. Dot plot of the best marker genes (d) as well as a heat map of the highest DEGs shows a unique expression profile for each microglia subtype. Scatter plot (f) and quantification (g) of flow cytometry analysis to validate the presence of DAM-A and its temporal changes after SCI. n=5 biological replicates for each time point in flow cytometry experiment.

We used flow cytometry to validate the presence of DAMs *in vivo* (gating strategy in Extended Data Fig 5). We focused on DAM-A due to its distinct molecular signature as compared to homeostatic microglia (Fig. 2), and its dramatic appearance and disappearance between 1dpi and 3dpi (Fig. 2b, c). Microglia were gated on P2ry12 and Msr1 expression based on our sequencing data (Fig. 2d). As expected, over 90% of microglia present in the uninjured spinal cord were in the homeostatic state (P2ry12^hi^/Msr1^lo^), and this decreased to 20% at 1dpi (Fig. 2f, g). At 1dpi, there was a large increase in Msr1^hi^ microglia, consistent with the appearance of DAM-A. However, the majority of Msr1^hi^ microglia were also P2ry12^hi^, representing a transition state between homeostatic and DAM-A, which are expected to be P2ry12^lo^/Msr1^hi^ based on our sequencing results. At 7dpi, we observed a significant decrease in Msr1^hi^ microglia and a partial return of homeostatic microglia, which this was not observed in our sequencing data. Taken together, our flow cytometry data support the appearance of DAM-A microglia subtype after SCI *in vivo*, but the temporal effects are more graded than those predicted from our sequencing data perhaps due to a delay in manifestation of gene expression changes at the protein level.

The peripherally-derived myeloid cluster revealed two monocyte and two macrophage subtypes in addition to border-associated macrophages and dendritic cells. The two monocyte subtypes corresponded to the classical Ly6C^hi^ and the non-classical Ly6C^lo^ monocytes (Fig. 3c) with the latter representing the largest population at 1dpi. 3 and 7dpi were dominated by the two macrophage subtypes, Macrophage-A and Macrophage-B respectively. Although these two subtypes did not correspond to the M1/M2 nomenclature^12^ (Extended Data Fig. 5), Macrophage-A expressed higher levels of the anti-inflammatory marker *arg1* than Macrophage-B (Fig. 3, Extended Data Fig. 5). This was consistent with Gene Ontology (GO) terms associated with inflammation being prevalent in Macrophage-B (Extended Data Fig 5). Both subtypes expressed the lysosomal gene *cd63*, but were distinguished by preferential expression of heme oxygenase *hmox in* Macrophage-A and *apoe* in Macrophage-B. In conclusion, our data reveals the presence of multiple cellular states in the monocyte-macrophage lineage that display temporal progression toward a more pro-inflammatory state. A putative minimal model depicting the relationship between these subtypes is illustrated in Extended Data Fig. 4b.

**Fig. 3.**
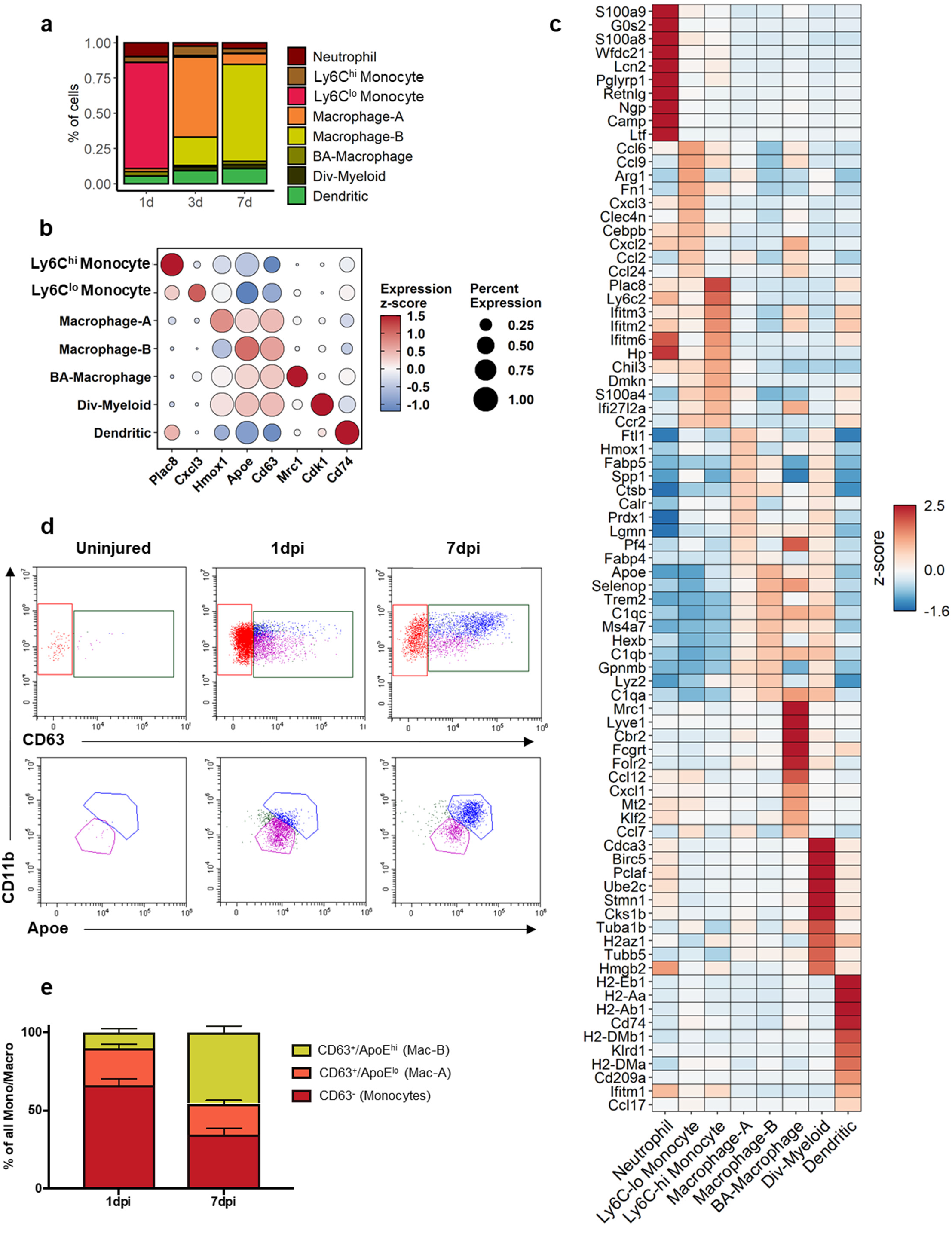
Molecular and temporal profile of monocyte/macrophage subtype heterogeneity acutely after SCI. (a) Percent of each peripheral myeloid subtype at each time point. (b) Dot plot of the best marker genes (c) as well as a heat map of the highest DEGs shows a unique expression profile for each peripheral myeloid subtype. Scatter plot (d) and quantification (e) of flow cytometry analysis to validate the presence of Macrophage-A and B and their temporal changes after SCI. n=5 biological replicates at each time point for flow cytometry experiment.

To validate the presence of Macrophage-A and Macrophage-B subtypes *in vivo*, we first isolated macrophages based on CD63^hi^ expression (gating strategy in Extended Data Fig. 6). Further separation on ApoE and CD11b expression revealed two distinct clusters that were consistent with Macrophage-A (CD63^hi^/ApoE^lo^/CD11b^med^) and Macrophage-B (CD63^hi^/ApoE^hi^/CD11b^hi^)subtypes (Fig. 3d, e). Consistent with our sequencing data, the monocytes were the predominant myeloid populations at 1 dpi, and subsequently decreased at 7dpi, whereas Macrophage-A (ApoE^lo^) and Macrophage-B (ApoE^hi^) were the most represented macrophage subtypes at 1 and 7dpi, respectively (Fig. 3e). Therefore, the flow cytometry data support the molecular identification of Macrophage-A and Macrophage-B subtypes and their temporal progression after SCI *in vivo*. In summary, analysis of myeloid cells reveal novel subtypes of microglia and macrophages during SCI progression.

### Vascular heterogeneity analysis identifies tip cell dynamics

To determine the heterogeneity of vascular cells, clustering analysis was performed only on the vascular clusters (endothelial cells, fibroblasts, and pericytes from Fig. 1) and visualized on a separate UMAP, which revealed endothelial cell subtypes on one side and perivascular mural cells on the other. We identified each cluster using annotated markers from a previous sc-RNAseq study of the brain vasculature^13^. We identified fibroblasts, pericytes, and vascular smooth muscle cells (VSMC) as three distinct populations of perivascular mural cells that were identified based on their expression of *col1a1, kcnj8*, and *acta2* respectively (Extended Data Fig. 7b). We also identified an unknown vascular subtype (U-Vascular) that clustered with mural cells due to its molecular similarity with pericytes, but also expressed endothelial cell markers (Extended Data Fig. 7c). Analysis of whether this could have been due to capture of two different cell types (i.e. doublet) during single cell processing indicated that this was not likely.

Next, we identified an arterial, a venous, and two capillary subtypes based on annotated markers^13^. Arterial endothelial cells were identified by expression of *gkn3* and *stmn2*, whereas venous endothelial cells were identified by *slc38a5* and *icam1*. Capillary endothelial cells were identified by the expression of general endothelial cell markers *ly6a* and *cldn5*, combined with the lack of selective arterial and venous markers (Fig. 4d, Extended Data Fig. 7c). The fifth endothelial cluster was identified as tip cells based on their expression of the canonical marker *apln*. Tip cells are leading cells present during vasculogenesis and angiogenesis, and they have not yet been systematically assessed in SCI. Whereas the other endothelial subtypes did not show large temporal changes in proportion, the proportion of tip cells increased significantly at 1dpi, and decreased to basal levels by 7dpi (Fig. 4c). This tip cell temporal profile was validated using *in situ* hybridization for *apln* combined with immunostaining for podocalyxin as an endothelial cell marker (Fig 4f). In the uninjured cord, *apln* transcripts were detected scattered throughout the gray matter, consistent with previous reports of *apln* expression in neurons^14^. However, we did not detect any endothelial cells expressing *apln* in uninjured tissue sections, suggesting that tip cells detected in uninjured tissue in our sc-RNAseq data was most likely due to tissue processing. Taken together, our analysis identified all known major vascular cell types at the injury site, including previously undescribed tip cell molecular and temporal profiles.

**Fig. 4.**
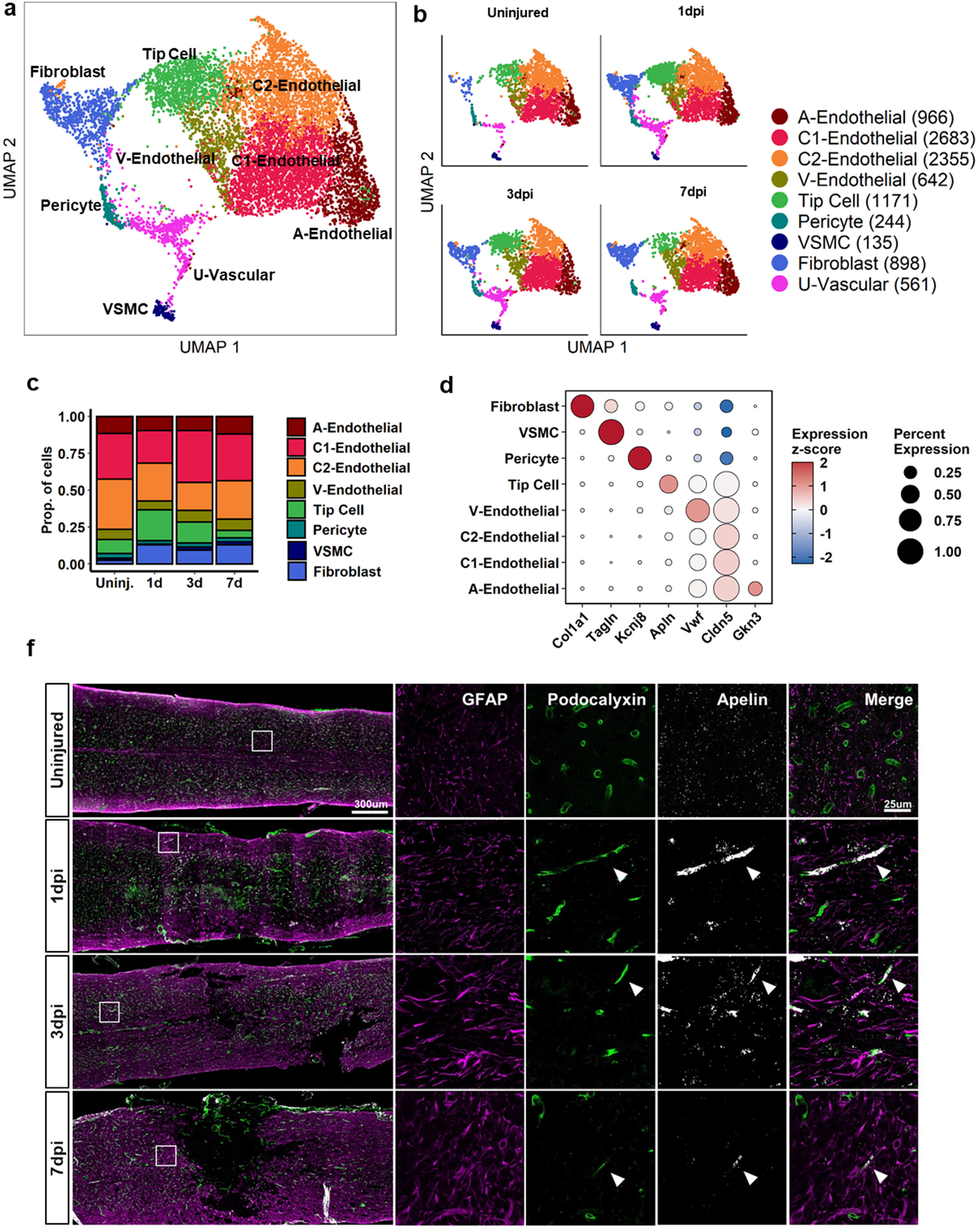
Molecular and temporal profile of vascular cell subtype heterogeneity acutely after SCI. (a) UMAP visualization of vascular cell clusters combined from uninjured, 1, 3, and 7dpi. Clusters were identified by previously annotated genes (Extended Data Fig. 7). (b) UMAP of each time point shows temporal progression of each cluster. (c) Quantification of the number and proportion of each cluster over time after injury. (d) Dot plot of the marker gene that best identifies each cell type. (f) Histological validation of the temporal progression of tip cells at the injury site. Uninjured, 1, 3, and 7dpi spinal cord sections were assessed by immunohistochemistry for podocalyxin (endothelial cells) combined with *in situ* hybridization for apelin (tip cells). GFAP was used as a marker for astrocytes to delineate the injury site. White box represents the corresponding region of the magnified images on the right. Arrow heads identify tip cells that co-express podocalyxin and apelin. Representative images from a total of 5 animals.

### Cellular interactions via angiopoietin and VEGF signaling during angiogenesis

To gain insight into mechanisms of cellular interactions during angiogenesis after SCI, we adapted CellPhoneDB^15^ to calculate “interaction scores” based on the average expression levels of a ligand and its receptor between two cells (Fig. 5a). Higher interaction scores indicate stronger predicted interactions between two cells via the specified ligand-receptor pathway. We focused on angiopoietin (Angpt) and vascular endothelial growth factor (Vegf) signaling due to their well-known roles in angiogenesis. Angpt1 is an agonist that promotes stabilization, whereas Angpt2 is an antagonist that promotes destabilization of blood endothelium by binding to the Tie2 receptor^16^. Endothelial subtypes did not show any significant differences in these ligand or receptor expression, and therefore were collapsed into a single endothelial category for simplification. The highest interaction score was detected between tip cells and endothelial cells along the Angpt2-Tie2 pathway at 1dpi (Fig. 5b). Expression analysis showed that Angpt2 is expressed most highly by tip cells at 1dpi followed by a gradual decrease over the next 7 days, and Tie2 (and Tie1) is expressed in endothelial cells at all time points (Fig. 5c). Pericytes and fibroblasts also contributed to this signaling interaction with endothelial cells, albeit at a lower level. These data suggest that Angpt2 has a dual role by autocrine signaling in tip cells that mediates their generation and behavior, as well as by paracrine signaling that destabilizes non-functional vessels. Interestingly, Angpt1 signaling to endothelial/tip cells shifted from VSMC at 1dpi to astrocytes at 3 and 7dpi (Fig. 5b). This corresponded to a shift in Angt1 expression from VSMC to astrocytes after injury (Fig. 5c), suggesting that astrocytes are important in vessel stabilization. Collectively, our data suggest that both vascular and neural cells interact with endothelial cells via Angpt signaling after SCI, with tip cells promoting vessel destabilization during early angiogenesis, and with astrocytes mediating vessel stabilization at later periods.

**Fig. 5.**
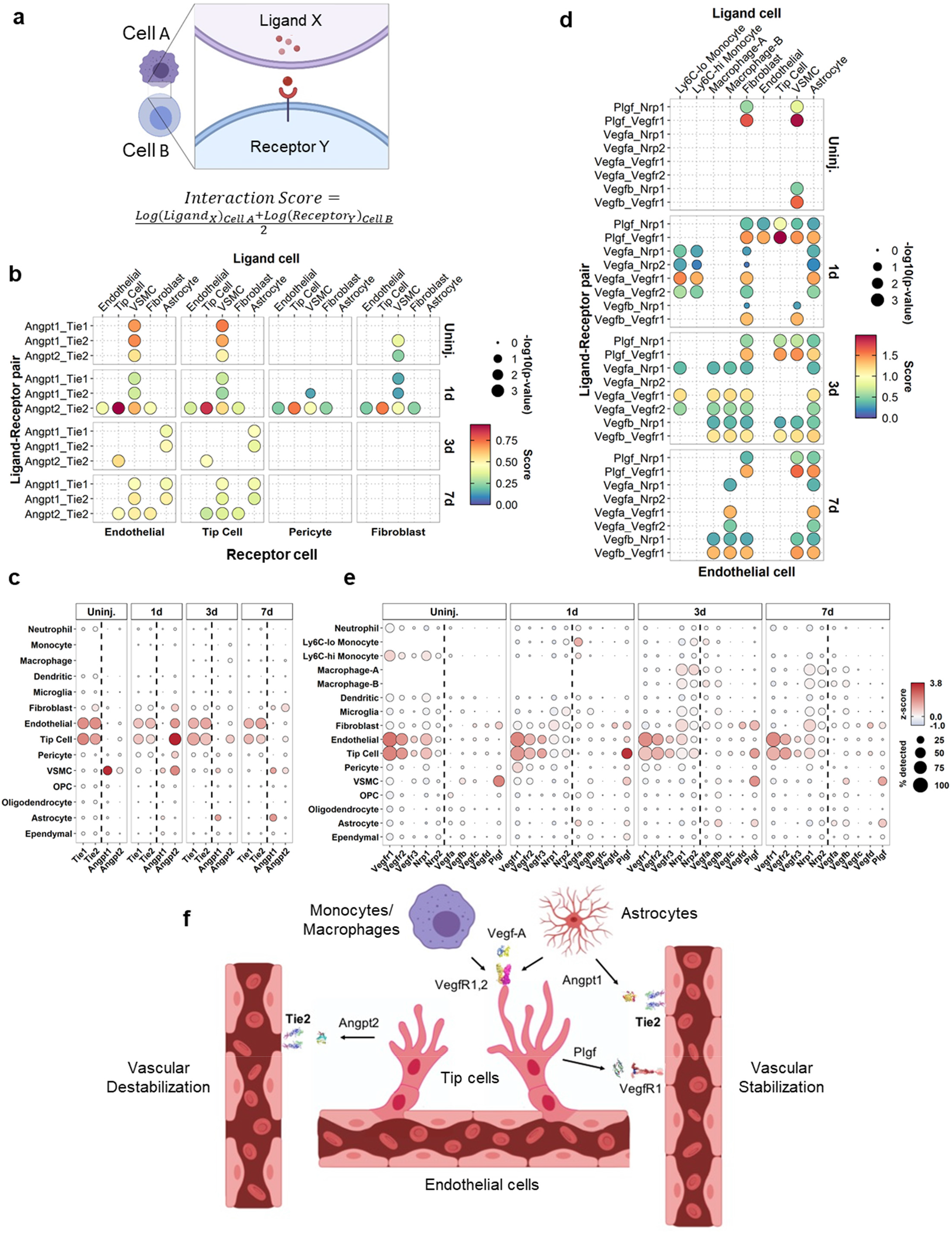
Ligand-receptor analysis of angiopoietin and vegf signaling reveals vascular interactions. (a) Interaction Score was calculated by the average of ligand expression in one cell type and receptor expression in another. Expression values were from a cell type cluster, and not from a single cell. (b) Dot plot of the interaction scores for angiopoietin signaling. Ligand expressing cells are listed on the top, and receptor expressing cells are on the bottom. The ligand-receptor pairs are listed on the left. Size of the circle denotes p-value, and color denotes interaction score. (c) Dot plot of expression values of angiopoietins and their receptors in each major cell type. (d) Dot plot of the interaction scores for vegf signaling. (e) Dot plot of expression values of certain vegf family members and their receptors in each major cell type. (f) Schematic depicting minimal model of vascular interactions via angiopoietin and vegf signaling during angiogenesis after SCI.

Vegfa binding to Vegfr1 and Vegfr2 on endothelial cells facilitates the proliferation, survival and directional sprouting of tip cell filipodia during angiogenesis^17–19^. The highest interaction scores for these ligand-receptor pairs were associated with monocytes/macrophages and astrocytes (Fig. 5d). Vegfa expression was highest in these two cell types at 1dpi, and Vegfr1 and Vegfr2 receptors were highly expressed by endothelial cells at all time points, suggesting that the major cues for new vessel formation is derived from the infiltrating myeloid cells and astrocytes (Fig. 5e). Strikingly, the strongest interactions amongst the Vegf family members were associated with placental growth factor (Plgf) binding to Vegfr1 (Fig. 5d). Plgf mediates angiogenesis by binding to Vegfr1^20, 21^, and our expression analysis showed tip cells to be the primary source, although lower levels of Plgf were also expressed by fibroblasts, VSMC, and astrocytes (Fig. 5e). Similar to our Angpt analysis above, our Vegf analysis suggests that tip cells play a major role in facilitating their own migration and growth, as well as those of other endothelial cells. Collectively, our data provide a rational and temporal framework of how endothelial cells interact with infiltrating myeloid cells and astrocytes via Angpt and Vegf pathways to re-vascularize the injured tissue, and highlight the autocrine and paracrine effects of tip cells in this process. A minimal model of revascularization in the first week after SCI is illustrated in Fig 5f.

### Analysis of macroglia heterogeneity reveals astrocyte and OPC-specific roles during gliosis

To assess macroglia heterogeneity, oligodendrocyte lineage cells, astrocytes, and ependymal cells were clustered and visualized on a separate UMAP (Fig. 6a, b), which showed spatial segregation corresponding to their major cell type. We identified three ependymal and an astroependymal subtypes with distinct expression and temporal profiles (Fig. 6c, e, Extended Data Fig 8). The astropendymal cells were best identified by their expression of the crystallin *crym*, and shared common markers with both astrocytes (e.g. *timp1, gfap*) and ependymal cells (e.g. *vim, tmsb10*). However, they did not express any cilia-associated genes that are hallmarks of ependymal cells (Extended Data Fig. 8c). The oligodendrocyte lineage cells segregated into previously described clusters that showed a spatial progression from OPC to dividing OPC and preoligodendrocytes, and finally to oligodendrocytes (Fig. 6a). These populations were identified by prototypical markers, with the exception of OPCs, which were best identified by tenascin *tnr* (Fig. 6e, Extended Data Fig. 8b). Whereas mature oligodendrocytes were predominant in the uninjured spinal cord, the proportion of OPCs, dividing OPCs, and preoligodendrocytes gradually increased over the next 7dpi. Taken together, our data highlight the heterogeneity of oligodendrocytes, astrocytes, and ependymal cells that reflect the effect of SCI on their differentiation state.

**Fig. 6.**
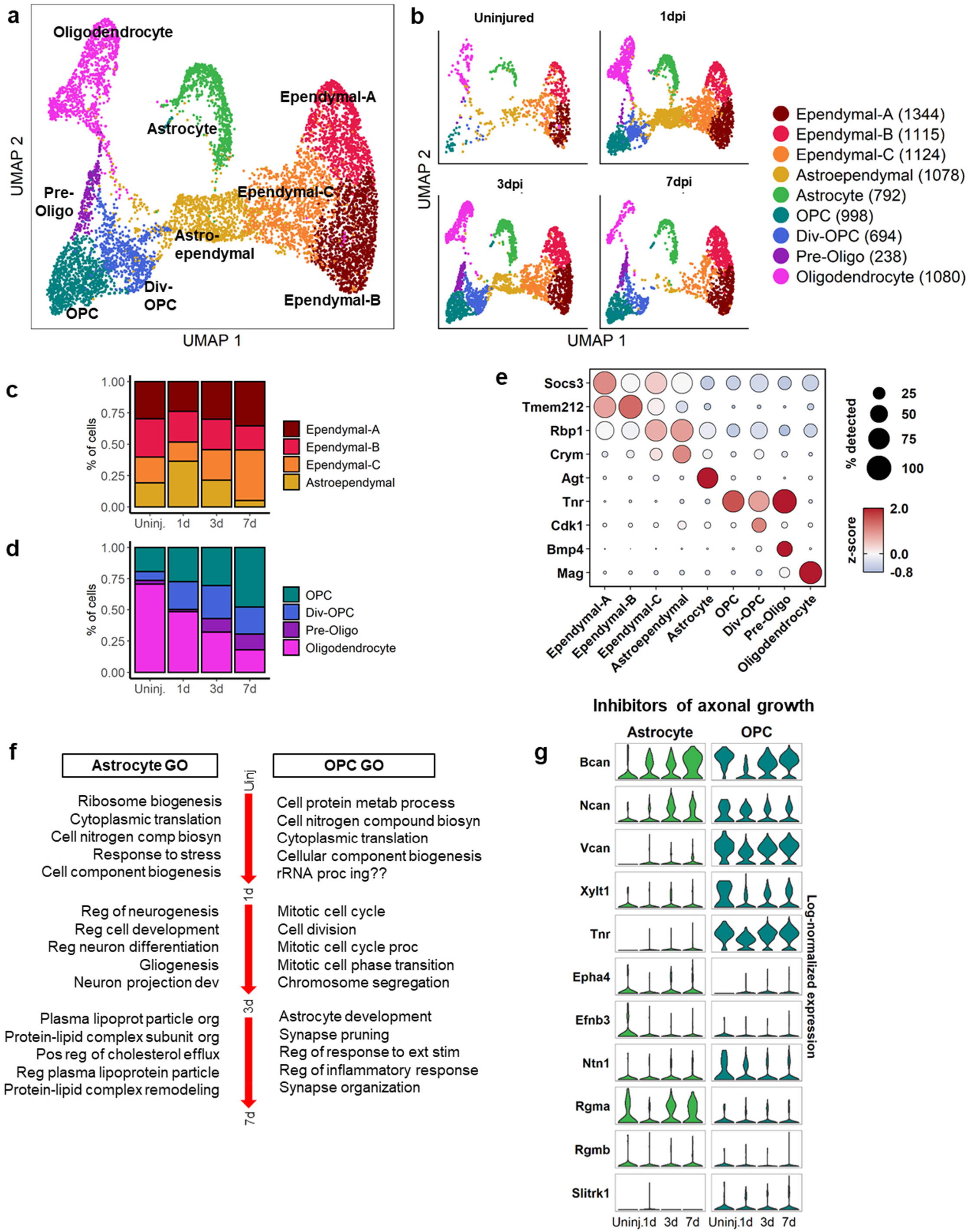
Molecular and temporal profile of macroglial heterogeneity acutely after SCI. (a) UMAP visualization of macroglial cell clusters combined from uninjured, 1, 3, and 7dpi. Clusters were identified by previously annotated genes (Extended Data Fig. 8). (b) UMAP of each time point shows temporal progression of each cluster. (c) Quantification of the number and proportion of each ependymal and (d) other macroglial clusters over time after injury. (e) Dot plot of the marker gene that best identifies each cell type. (f) Comparison between astrocyte and OPC Gene Ontology biological processes based on DEG between sequential time points. GO terms are listed in rank order from top to bottom for each time point. Remaining GO terms are in Expanded Data Fig. 9. (g) Comparison of expression of common axon growth inhibitors between astrocytes and OPCs.

Although gliosis has been synonymous with reactive astrocytes, accumulating evidence indicates that OPCs are also an important component of the glial scar^22, 23^. To compare reactive OPCs and astrocytes, we performed Gene Ontology (GO) Enrichment Analysis for biological processes associated with DEGs between each time point (Fig. 6f). At 1dpi, top biological processes for both astrocytes and OPCs pertain to translation and biogenesis. By 3dpi, astrocytes are defined by processes related to neurogenesis and gliogenesis, whereas OPCs are defined by mitosis, which reflect the active state of proliferation and differentiation for both cell types during this time. Interestingly, the DEGs between 3 and 7dpi in astrocytes were related to lipid processing, whereas those in OPCs were related to several processes including astrocyte development, synapse remodeling, and the inflammatory response, which are processes that have been typically associated with reactive astrocytes. To assess the potential effects of reactive astrocytes and OPCs on axonal growth, we compared the expression levels of axon growth inhibitory molecules using a previously curated list^24^ (Fig. 6g). Interestingly, inhibitory proteoglycans such as *acan, bcan, ncan*, and *vcan* were expressed preferentially by OPCs. In summary, our data suggest that although astrocytes and OPCs initially display similar generic responses to injury, by 7dpi, they acquire unique functional identities, and many processes that have traditionally been attributed to reactive astrocytes are also present in OPCs.

### Macrophage-Mediated Mechanisms of Gliosis and Fibrosis

Previous studies have shown that STAT3 plays an important role in astrogliosis^4, 25, 26^, but its upstream activators have yet to be clearly defined. Since IL6 cytokine family members are the main STAT3 activators, we assessed their expression across all cell types and found oncostatin M (*osm*) expressed at highest levels in myeloid cells, *il6* expressed at highest levels in fibroblasts, and *clcf1* expressed at highest levels in astroependymal cells (Fig. 7a). Other IL6 cytokines were either not expressed highly or dropped out of our sequencing analysis. Oncostatin M receptor (*osmr*) and the signaling coreceptor gp130 (*il6st*) were expressed highly in fibroblasts and astrocytes, and interaction scores for IL-6 cytokine family members were highest for OSM signaling between Ly6C^lo^ monocytes/Macrophage-B and astrocytes/fibroblasts (Fig. 7b). To validate the sequencing data, we performed double *in situ* hybridization for *osm* and *cd11b* and found a significant increase in *osm* mRNA in *cd11b^+^* myeloid cells compared to non-myeloid cells (Fig. 7c, e). To assess OSMR expression, we used immunohistochemistry and found increasing expression of OSMR in both Pdgfr-β^+^ fibroblasts and GFAP^+^ astrocytes in the fibrotic and glial scar, respectively (Fig. 7d, f, g). Taken together, our results strongly suggest that the gp130 signaling pathway induced by Osm is a common mechanism by which astrocytes and fibroblasts are preferentially activated by specific monocyte/macrophage subtypes after SCI.

**Fig. 7.**
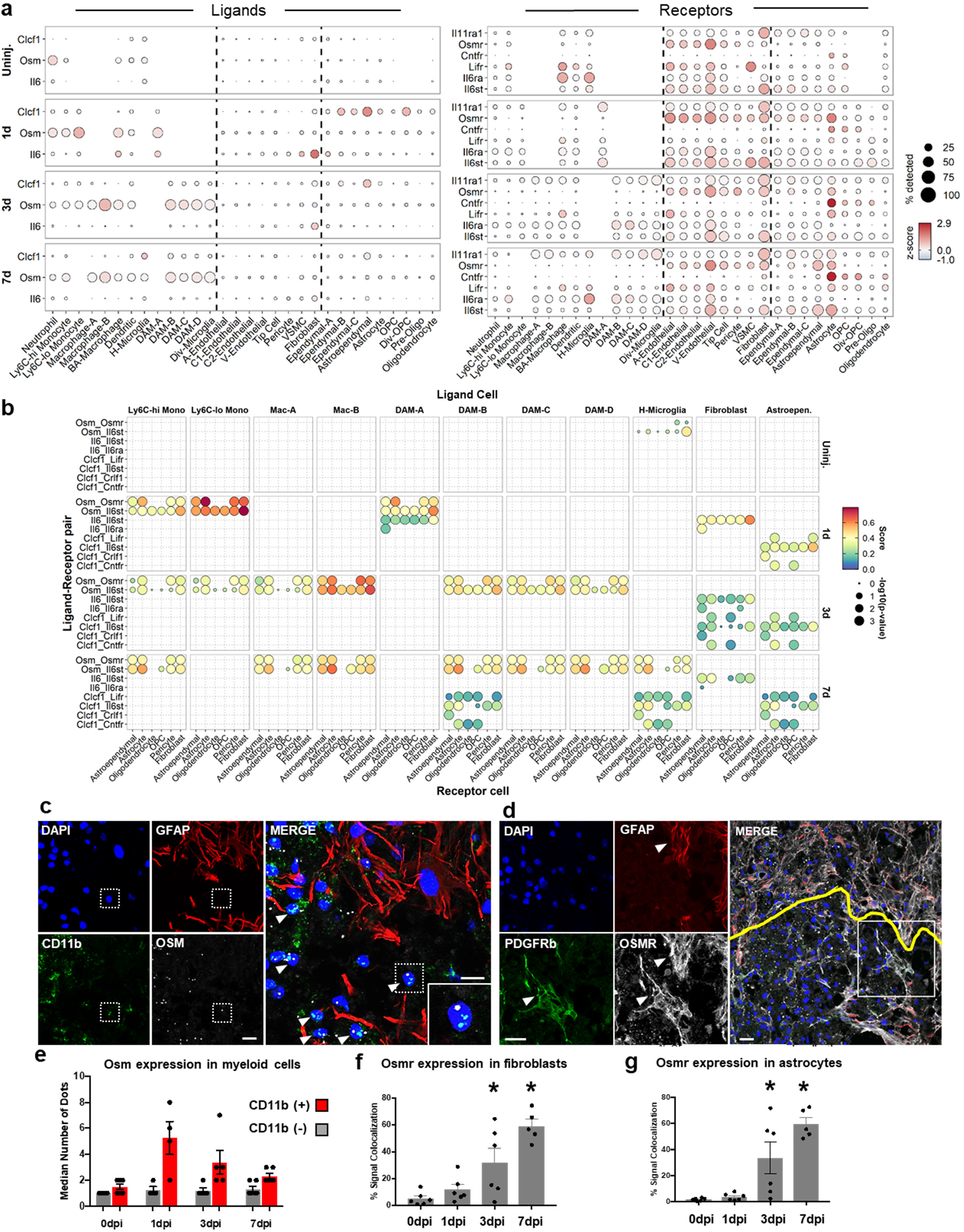
Ligand-receptor analysis of IL6 cytokine family members identify Oncostatin M signaling between myeloid cells and astrocytes/fibroblasts. (a) Dot plot depicting expression levels of IL6 family member ligands and receptors. The size of the dot depicts the percent of cells expressing the gene at the given time. Ligands/receptors not on the list did not reach statistical significance or were not present in the sequencing data. (b) Dot plot of interaction scores of IL6 family member signaling show highest predicted interaction between myeloid cells and astrocytes/fibroblasts via oncostatin M signaling. (c) Histological validation of oncostatin M expression in myeloid cells. Spinal cord sections were assessed by immunohistochemistry for GFAP (astrocytes) combined with *in situ* hybridization for CD11b (myeloid) and Osm (oncostain M). DAPI denotes cell nuclei. (d) Histological validation of oncostatin M receptor (Osmr) expression in astrocytes and fibroblasts. Spinal cord sections were assessed by immunohistochemistry for GFAP (astrocytes), Pdgfr-β (fibroblasts), and Osmr. Yellow line delineates astroglial scar border. (e) Quantification of Osm expression in myeloid (DAPI^+^/CD11b^+^/Osm^+^) and non-myeloid (DAPI^+^/CD11b^−^/Osm^+^) cells shows significantly higher Osm expression in myeloid cells. Quantification showed increasing colocalization of Osmr with Pdgfr-β (f) or GFAP (g) immunoreactivity over time. Error bars represent SEM. *p<0.05 compared to 0dpi. One-Way ANOVA and Tukey’s post-hoc test. Images are from 7dpi. Scale bar = 15μm. Each data point represents a biological replicate.

To identify other distinct and common pathways by which macrophage subtypes mediate astrogliosis and fibrosis, we calculated interaction scores for all known ligand-receptor pairs between Macrophage-A/B and astrocytes/fibroblasts at 3 and 7dpi (Fig. 8). While many ligands expressed by macrophages signaled to receptors on both astrocytes and fibroblasts (i.e. common pathways), there were many more ligand-receptor pairs unique to macrophage-fibroblast interactions than macrophage-astrocyte interactions (i.e. distinct pathways). These unique macrophage-fibroblast interactions included signaling related to IL1α/β, Vegfa/b, Pdgfα, and Tgfβ1. Overall, the highest interactions scores were associated with Spp1 and Apoe signaling, which were common to both astrocytes and fibroblasts. Macrophages also displayed subtype specificity in signaling to astrocytes and fibroblasts. For example, Jag1-Notch signaling was largely specific to Macrophage-A-astrocyte interaction, whereas IL1α/β-IL1r1 signaling was specific to Macrophage-B-fibroblast interaction. In summary, our analysis highlights the utility of our sc-RNAseq dataset in identifying potential signaling mechanisms that mediate astrogliosis and fibrosis after SCI.

**Fig. 8.**
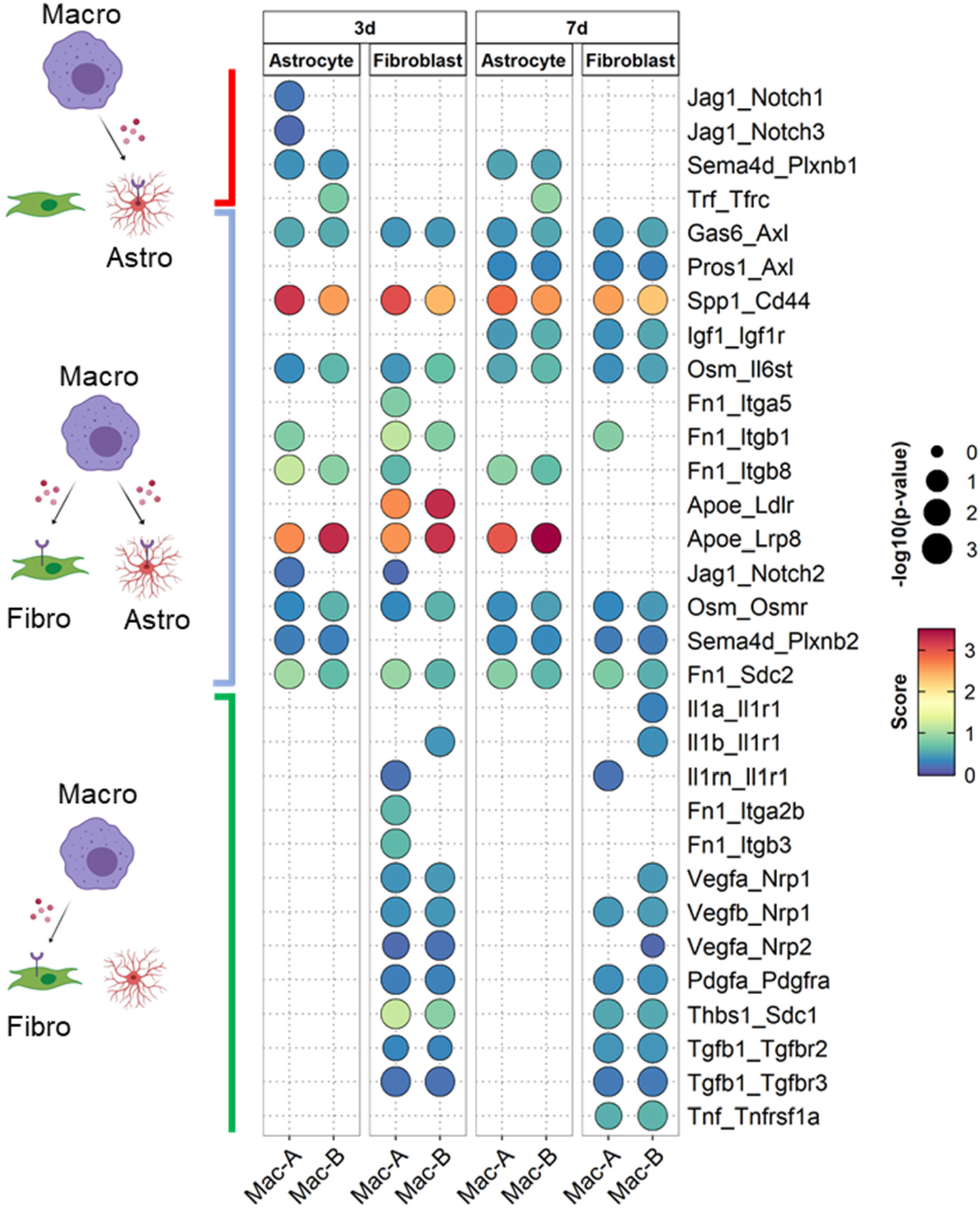
Macrophage subtypes interact with astrocytes and fibroblasts via distinct and common signaling pathways. Dot plot of interaction scores for all ligand-receptor pairs at 3 and 7dpi showed signaling specific between macrophages and astrocytes (red bracket on top) as well as between macrophages and fibroblasts (green bracket on bottom). Common pathways are indicated by the blue bracket in the middle. Macrophage subtypes (A or B) are denoted at the bottom.

## Discussion

The overall goal of this study was to use sc-RNAseq to generate a resource with which we can gain insight into the complex cellular heterogeneity and interactions that occur at the spinal cord injury site. We were successful in isolating virtually every major cell type known to be present at the injury site, which provided an opportunity to assess the wound healing process with much less bias than prior studies that focused on single cell types. In this regard, we found that myeloid subtypes display distinct temporal regulation of angiogenesis, gliosis, and fibrosis. These results support previous studies showing that myeloid depletion leads to reduced angiogenesis, astrogliosis, and fibrosis after CNS injury^3, 27, 28^, and provide further insight by identifying potential contributions of specific myeloid subtypes.

We can begin to infer putative functions of the two macrophage subtypes by placing their temporal progression in context of prior studies. Our finding that Macrophage-A and B subtypes do not correspond to the M1/M2 nomenclature (Extended Data Fig. 5) is consistent with findings from other sc-RNAseq studies^29–31^. However, there are parallels with the generally accepted paradigm that macrophages shift from an anti-inflammatory to a more pro-inflammatory state over time after SCI^32–34^. As the major peripheral myeloid composition at the injury site shifts from monocytes to Macrophage-A and then to Macrophage-B over time, there is a progressive decrease in the classic anti-inflammatory enzyme Arginase 1 that is associated with increased pro-inflammatory IL1 and TNF pathways in Macrophage-B. Despite these pro-inflammatory functions, our data identified several mechanisms by which macrophages may have beneficial effects on the wound healing process. For example, Macrophage-B express Vegfa that is well-known to be necessary for growth and differentiation of endothelial cells during angiogenesis^35^. In addition, macrophages are a major source of osteopontin (*spp1*) and ApoE, both of which have neuroprotective roles after SCI. Spp1 and ApoE knockout mice both display worse histopathology and behavioral recovery after SCI, whereas administration of ApoE mimetic improves recovery^36–38^. Thus, a future challenge is to limit the pro-inflammatory macrophage state without interfering with their beneficial effects on wound healing.

Of the microglial subtypes, DAM-D is notable for its expression profile defined by neuroprotective/neuroregenerative genes. In addition to osteopontin and ApoE whose protective roles were mentioned above, the well-known neuroprotective growth factor Igf-1^39^ was the best marker gene for DAM-D. Interestingly, a recent microglia depletion study demonstrated that microglia have a neuroprotective role by inducing astrogliosis via Igf-1^28^. This study also reported that scar-forming microglia that are located between the astroglial and fibrotic scars are the major source of Igf-1 at 7dpi, which is consistent with the appearance of DAM-D in our data, and suggests a spatial specificity for DAM-D. In addition, promoting proliferation of microglia reduced the lesion area and improved behavioral recovery, which supports the working model of dividing microglia giving rise to DAM-D. In addition to their neuroprotective effects, Igf-1 and osteopontin have been shown to promote axonal regeneration after SCI^40–42^, which make DAM-D an attractive therapeutic target for neurological disorders.

The vascular analysis revealed novel insight in the contribution of tip cells and astrocytes during angiogenesis after SCI. The data indicate that tip cells are highly dynamic; they appear quickly at 1dpi and are mostly gone by 7pi. They are the major source of Angiopoietin-2, which destabilizes vessels, as well as placental growth factor, which promotes vessel formation, suggesting that tip cells mediate vessel remodeling during the few days after SCI. Interestingly, their disappearance overlaps with increased expression of Angiopoietin-1 in astrocytes. Angiopoietin-1 typically promotes vessel stabilization, suggesting that astrocytes contribute to vessel maturation after the initial phase of vascular remodeling by tip cells. This is corroborated by previous study demonstrating that when the blood-spinal cord barrier is restored by 21dpi, blood vessels closely associated with astrocytes in the glial scar region express Glut1, whereas vessels in the astrocyte-devoid fibrotic region do not express this mature blood-spinal cord barrier marker^5^. These data highlight an understudied role of astrogliosis in vessel maturation after SCI, and provide new insight into prior astrocyte ablation studies that suggested “corralling” of infiltrating leukocytes by the astroglial scar^25, 43^. Our data raise the possibility that reducing astrogliosis prevents maturation of blood vessels, which leads to increased infiltration of leukocytes into the spinal cord parenchyma surrounding the injury site. Other possible mechanisms of interactions between astrocytes, vasculature, and axons need to be explored in the future.

Our single cell RNA-seq dataset is the first comprehensive transcriptomic analysis that captures virtually all cells that contribute to the injury site pathology after SCI. This dataset can be used to assess not only heterogeneity of the cells that comprise the injury site, but also to assess signaling mechanisms that underlie cellular interactions at the injury site. Our own analysis revealed novel insight into myeloid cell heterogeneity, and specific signaling pathways by which unique myeloid subtypes contribute to the wound healing process, including angiogenesis and scar formation. These new insights can help decode the complex processes that underlie spinal cord injury pathobiology.

## Acknowledgements

Authors would like to thank the SCCC Onco-Genomics Shared Resource Core, Division of Veterinary Resources, DRI Flow Cytometry Core, HIHG Bioinformatics Core, Yan Shi at the Miami Project to Cure Paralysis Microscopy Core. Special thanks to Shaffiat Karmally and Yadira Salgueiro for their assistance in multiple aspects of this study, including animal care and logistics. This study was funded by NINDS R01NS081040 (J.K.L), University of Miami SAC Award UM SIP 2019-2 (J.K.L), NINDS F31NS115225 (C.B.R), the Miami Project to Cure Paralysis and The Buoniconti Fund, a generous gift from the Craig H. Nielsen Foundation, and NIH 1S10OD023579-01 for the VS120 Slide Scanner housed at the University of Miami Miller School of Medicine Analytical Imaging Core Facility.

## Author Contributions

L.M., J.S.C., J.K.L, C.B.R. designed and performed the experiments, analyzed the data, and wrote the manuscript. S.L.Y. performed the experiments and wrote the manuscript. P.T. designed the experiments, analyzed the data, and wrote the manuscript.

## Competing Interests

The authors declare no competing interests.

## Methods

### Mice & Spinal Cord Injuries

For sequencing, flow cytometry, and histological validation of tip cells, C57BL/6J mice 8-10 weeks of age were purchased from Jackson Labs (Stock 000664). In total, we sequenced 2 biological replicates for each of the four time points to yield 8 sample datasets. For the first replicate at each time point, mice received spinal cord injury or no injury as described below and injury site (or corresponding region in uninjured controls) from each mice were individually processed for single cell RNA-seq (scRNA-seq). For the second replicate at each time point, we pooled cells from 2 animals using a modified tissue dissociation procedure as described below. For histological validation of activated fibroblasts, Postn-CreER mice (Jackson Laboratory stock 029645) were bred to Rosa26-EYFP mice (Jackson Laboratory stock 006148) to generate Postn^EYFP^ mice. All mice were of C57BL/6 background. To induce EYFP expression, Postn^EYFP^ mice were injected i.p (intraperitoneal) with 0.124mg/g body weight of tamoxifen (MP Biomedicals, dissolved in 90% sunflower oil, 10% ethanol) for 5 consecutive days. Contusive spinal cord injuries were performed 7 days after the last tamoxifen injection.

Contusive spinal cord injuries for all mice were performed as previously described^3^. Briefly, mice were anesthetized using ketamine/xylazine (100mg/15mg/kg i.p.) and received a 65 kdyne mid-thoracic (T8) contusion injury. Laminectomies were performed at the T8 vertebral level followed by stabilization via a spinal frame clamping the T7/T9 spinous processes. A computer controlled contusion injury was delivered using the Infinite Horizons Impactor device (Precision Systems and Instrumentation, LLC). Injured mice were given post-operative fluids subcutaneously consisting of Lactated Ringer’s solution (1mL), gentamycin (5 mg/kg) and buprenorphine (0.05 mg/kg) twice daily for 7 days. Bladders were manually expressed twice daily until the end of the experiment. All procedures were in accordance with University of Miami IACUC and NIH guidelines.

### Tissue Dissociation

Mice were anesthetized with Avertin (250mg/kg, i.p) before transcardial perfusion with artificial CSF solution (87mM NaCl, 2.5mM KCl, 1.25mM NaH2PO_4_, 26mM NaHCO_3_, 75mM sucrose, 20mM glucose, 1.0mM CaCl_2_, 2mM MgSO_4_). Solution was oxygenated on ice for 10min before use. An 8mm section of spinal cord centered at the injury site (or the corresponding location in uninjured tissue) was dissected. After removal of the meninges, tissue was chopped using a razor blade, washed with 5mL CSF solution, and centrifuged at 300g for 5min at 4°C. The pelleted cells were processed using the Miltenyi Neural Tissue Dissociation Kit (P) (Cat #: 130-092-628) following manufacturer’s suggested protocol to obtain a single cell suspension. In brief, the pellets were incubated in a 2mL of Enzyme Mixture 1 for 30min at 37°C (50μL Enzyme P in 1.9mL Buffer X) with gentle shaking by hand every 5 minutes. 30μL of Enzyme Mixture 2 (10 μL Enzyme A in 20 μL Buffer Y) was added to the cell suspensions and manually triturated (slowly, 10 times per sample) with a large (1000μm diameter) opening fire-polished pipette. After 10min incubation at 37°C, suspensions were triturated with a medium and then small (750μm and 500μm diameter respectively) opening fire-polished pipette to produce single cell suspensions. Suspensions were strained through a 70μM cell strainer and washed with 10mL CSF solution. Suspensions were centrifuged at 300g for 5min and supernatants were discarded. Cell pellets were incubated in 10μl Miltenyi Myelin Removal Beads II (Cat #: 130-096-733) and 90μL MACS Buffer (0.5% BSA in HBSS w/o Ca2+/Mg2+) for 15min at 4°C. Cells were washed with 10mL MACS Buffer and centrifuged at 300g for 5min. Supernatants were discarded and pellets were resuspended in 1 mL MACS buffer. Suspensions were then applied onto equilibrated LS Miltenyi MACS Magnetic Bead Columns (Cat #: 130-042-401). Columns were washed with 2mL MACS buffer, flow through was centrifuged at 300g for 5min, and supernatants were discarded.

Due to the low number of astrocytes recovered in our first set of replicates, we enriched our second set of replicates with astrocytes by pooling from two spinal cord samples. We processed two spinal cords from each injury time point in parallel as described above to yield two suspensions per time point. One suspension from each time point was kept on ice while the was further processed with the Miltenyi Anti-ACSA-2 MicroBead kit according to manufacturer’s instructions. In brief, suspensions were incubated with Miltenyi ACSA2 Beads for 15mins at 4°C and subsequently washed with 1mL of MACS buffer. Suspensions were applied to LS Miltenyi MACS Magnetic Bead Columns and remaining cells were flushed from each column using LS Miltenyi column syringes and collected in separate tubes. ACSA-2 enriched flow throughs were then pooled with the single cell suspension from the other tissue.

Combined suspensions were centrifuged at 300g for 5min and the pellets were resuspended in Red Blood Cell Lysis buffer (155 mM NH_4_Cl, 10 mM KHCO_3_, 0.09 mM Na4-EDTA) and incubated at room temperature for 1min. After washing with 5mL MACS Buffer, suspensions were centrifuged at 300 g for 5 min and supernatants were discarded. Pellets were resuspended in 50μl FACS buffer (1% BSA + 0.05% sodium azide in HBSS w/o Ca2+/Mg2+) and processed for scRNA-seq. Before submitting for sequencing, each sample underwent a quality control assessment which included cell viability and number. Briefly, cell suspensions were incubated with ViaStain AOPI Staining Solution and cell concentration/viability was assessed using the Nexcelom Cellometer K2. Each submitted sample met a viability cutoff of 80% or higher. Each sample was diluted accordingly to achieve a target cell capture of 10,000 cells, with our actual capture for the two replicates totaling 4721 for Uninjured, 14536 for 1dpi, 19405 for 3dpi, and 18446 for 7dpi.

### Histology

Mice were anesthetized with Avertin as described above and transcardially perfused with cold 4% paraformaldehyde (PFA) in PBS. Spinal cord was removed, and following 2 hours of post-fixation in 4% PFA, it was placed in 30% sucrose (in PBS) solution overnight on a shaker. 8mm segment of the spinal cord centered at the injury site (or the corresponding region in uninjured spinal cord) was embedded in OCT compound (Tissue-Tek) and sectioned sagittally on a cryostat (16μm). Serial tissue sections were mounted onto Superfrost Plus slides, and was stored at −20°C.

#### Immunohistochemistry

After drying at room temperature, tissue sections were washed once with PBS, followed by permeabilization with PBS-T for 10 min at RT (0.3% Triton X-100 in PBS). After 1hr of incubation in blocking solution at room temperature (5% normal donkey serum in PBS-T), sections were incubated in primary antibody diluted in 5% normal donkey serum overnight at 4°C (primary antibodies are listed in the table below). After primary antibody incubation, slides were washed 2 times with PBS for 2 min each. Slides were incubated with appropriate AlexaFluor secondary antibodies (1:500 in PBS-T) for 1 hr at room temperature. Slides were washed with 2 times PBS for 2 min each and incubated with DAPI (1:15,000) for 3min at room temperature in the dark. Slides were washed with PBS and glass coverslips were mounted with Fluoromount-G (Southern Biotech Cat #: 0100-01). Images were taken on an Olympus Fluoview1000 Confocal Microscope or on an Olympus VSI Slide Scanner.

| Antibody | Company | Catalog #/RRID | Dilution |
| --- | --- | --- | --- |
| Chicken anti-GFAP | Abcam | ab4674/AB-304558 | 1:500 |
| Rat anti-GFAP | Life Technologies | 13-0300/AB-86543 | 1:500 |
| Rabbit anti-PDGFR (Y92) | Abcam | ab32570/AB-777165 | 1:200 |
| Goat anti-OSMR | R&D Systems | AF-662-SP/AB-355511 | 1:100 |
| Goat anti-Podocalyxin | R&D Systems | AF1556/AB-354858 | 1:200 |

#### In situ hybridization

Tissue sections were prepared as described above. *In situ* hybridization was performed using the RNAscope Multiplex Fluorescent Kit v2 (Advanced Cell Diagnostics Cat #: 323100) with a few modifications to the manufacturer’s protocol. All incubations were at room temperature unless otherwise noted. Briefly, slides were incubated in hydrogen peroxide for 10min. After washing with distilled water, slides were incubated in 1X Target Retrieval Solution at 95°C for 1min. After washing with distilled water, slides were incubated in 100% EtOH for 3min. Slides were left to dry before incubation in RNAscope Protease III solution at 40°C for 30min (all 40°C incubations were in the Advanced Cell Diagnostics HybEZ Oven). After incubation, slides were washed with distilled water and incubated in probe solution for 2hrs at 40°C. The following probes were purchased from Advanced Cell Diagnostics: C2 Mm-OSM (Cat #: 427071-C2 (1:50)), C1 Mm-ITGAM (Cat #: 311491 (no dilution, used working solution)), C2 Mm-Apln (Cat #: 415371-C2 (1:50)). Following probe hybridization, each amplifier was individually incubated with slides for 30min at 40°C. Before each fluorophore incubation, slides were incubated in RNAscope HRP solution for 15min at 40°C. Fluorophores for each channel were prepared at 1:750 in RNAscope Diluent Solution, and individually incubated with slides for 30min at 40°C. Different combinations of TSA Plus Cyanine 5 (Perkin Elmer Cat #: NEL745001KT) and TSA Plus Fluorescein (Perkin Elmer Cat #: NEL74100KT) were used with different probe channels. After each individual fluorophore incubation, slides were washed with RNAscope 1X Wash Buffer and incubated with RNAscope HRP Blocker solution for 15min at 40°C. After *in situ* hybridization, slides were washed with PBS and immunohistochemically stained with appropriate primary antibodies as described above.

### Tissue Quantifications

For each sagittal tissue section of the OSMR/GFAP/PDGFR-β and OSM/GFAP/CD11b stain (immunohistochemistry and in situ hybridization, respectively), three 40X confocal images were taken surrounding the injury site (3 serial tissue sections were analyzed per animal). Images were analyzed using the Cellomics High-Content Screening software. To quantify total signal colocalization between GFAP/PDGFR-β and OSMR, we used the Target Activation V3 algorithm, with a minimum intensity threshold set to 350 for OSMR fluorescence. To count the number of OSM+/CD11b+ nuclei within the injury site, we used the Colocalization V4 algorithm.

### Flow Cytometry

Mice received spinal cord injury as described above, and injury sites were processed for flow cytometry. Mice were anesthetized with Avertin followed by transcardial perfusion with ice-cold PBS. An 8mm segment of the spinal cord centered at the injury site was dissected and the meningeal layer was removed. Spinal cords were manually dissociated by mechanical grinding against a 70μm cell strainer (the plunger of a 3mL syringe was used for mechanical dissociation). Strainers were washed with 10mL HBSS (w/o Ca2+/Mg2+) and centrifuged at 300g for 10min. Supernatant was aspirated and the pellet was resuspended in 10μL Miltenyi Myelin Removal Beads II (Miltenyi Biotec Cat #: 130-096-733) and 90 μL MACS Buffer (0.5% BSA in HBSS w/o Ca2+/Mg2+) and incubated for 15min at 4°C. Cells were washed with 10mL MACS Buffer and centrifuged at 300g for 10min. Pellet was resuspended in 1mL MACS buffer and applied onto equilibrated LS Miltenyi MACS Magnetic Bead Columns (Miltenyi Biotec Cat #: 130-042-401). Columns were washed with 2mL MACS buffer and suspension was centrifuged at 300g for 5min. Pellet was resuspended in RBC Lysis buffer (155mM NH_4_Cl, 10mM KHCO_3_, 0.09mM Na4-EDTA) and kept at room temperature for 1min. To identify live cells, pellet was resuspended in Zombie Aqua Viability Dye and incubated for 30min at room temperature (Biolegend Cat #: 423101). Following a wash in PBS and spin at 300g, cells were resuspended in Fix/Perm Solution for 20min at 4°C to fix and permeabilize the cells (BD Bioscience Cytofix/Cytoperm Cat #: 554714). After incubation, cells were washed with 1mL of Perm/Wash solution. After centrifugation at 300g for 10min, supernatant was aspirated and the cells were incubated in blocking buffer mixture for 5min at 4°C (1uL anti-mouse CD16/32 TruStain Fcx (RRID: AB-1877271 AB-1574973, Biolegend Cat #: 101320) and 99uL FACS buffer (1% BSA + 0.05% sodium azide in HBSS w/o Ca2+/Mg2+)). Antibody mixture was added directly to the cell suspension and incubated for 20min at 4°C. Cells were washed with 5mL FACS buffer and centrifuged at 300g for 10min. For fixation, cells were resuspended in 4% PFA and incubated for 30min at 4°C. After centrifugation at 300g for 10 min, cells were resuspended in 250μL FACS buffer and transferred into a flat bottom 96-well plate. 10μL of 123 eBeads Counting Beads (ThermoFischer Scientific Cat #: 01-1234-42) were added to each sample well and cells were kept at 4°C until analysis on a Cytoflex Flow Cytometer (Beckman Coulter).

#### Macrophage Flow Antibody Panel

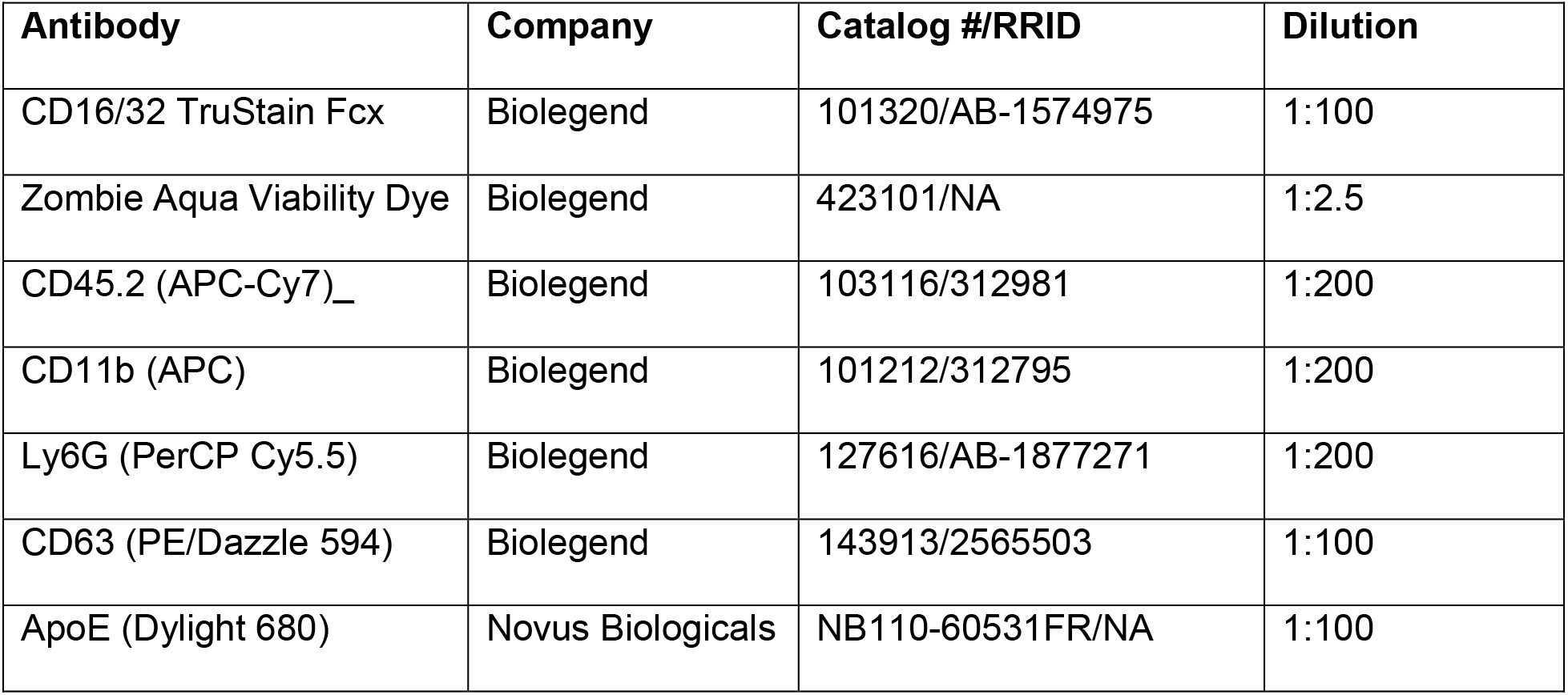

#### Microglia Flow Antibody Panel

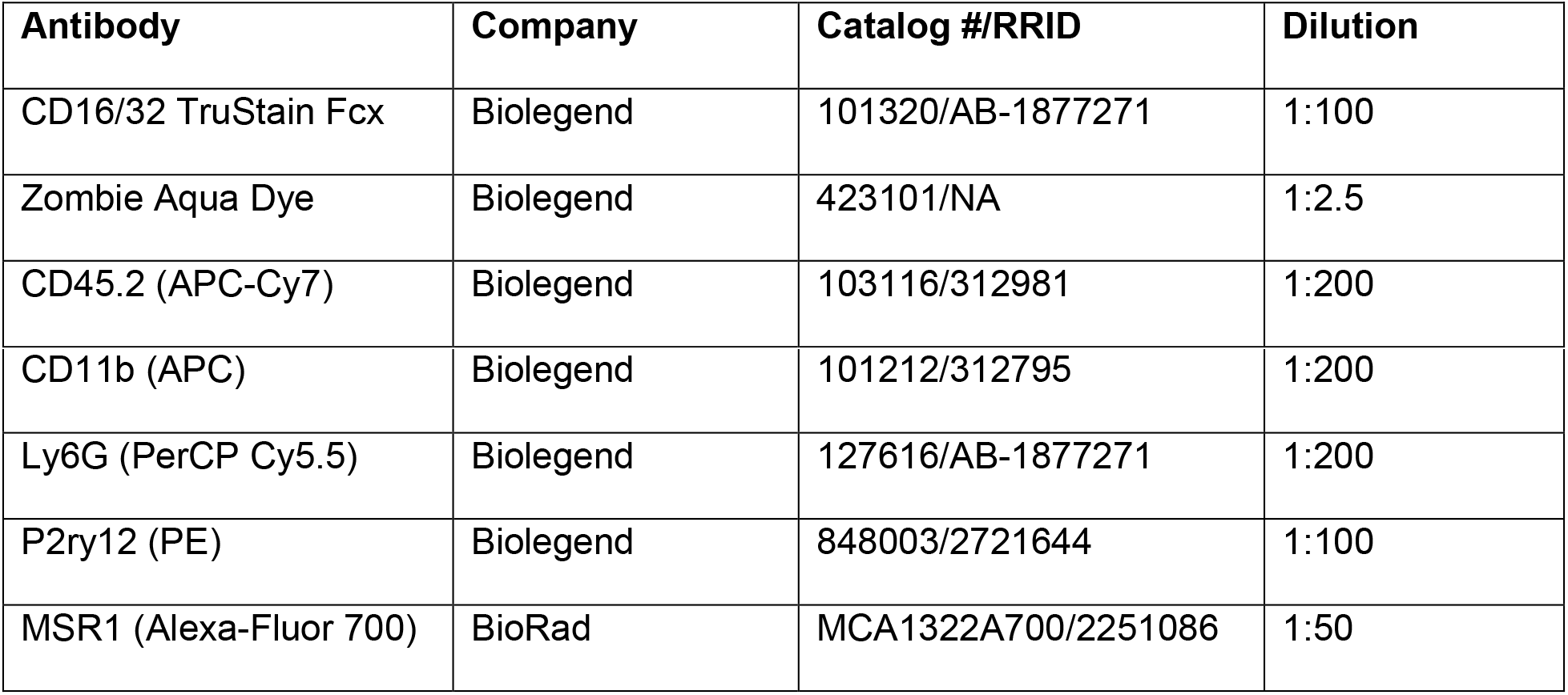

### Single Cell Sequencing & Bioinformatics

| Software, Algorithms, and Data | Source | Version |
| --- | --- | --- |
| CellRanger | 10X Genomics | 2.2.0 and 3.0.1 |
| RStudio | Rstudio, Inc. | 1.2.5001 |
| R | The R Foundation | 3.6.1 |
| Seurat | https://satijalab.org/seurat/ | 3.1.2 |
| ggplot2 | Wickham H (2016)<sup>44</sup> | 3.3.0 |
| dplyr | Wickham H et al. (2020) | 0.8.5 |
| scater | McCarthy et al (2017)<sup>45</sup> | 1.12.2 |
| DropletUtils | Lun et al. (2019)<sup>46</sup> | 1.4.3 |
| python | Python Software Foundation | 3.7.6 |
| Scrublet | Wolock et al. (2019)<sup>47</sup> | 0.2.1 |
| topGO | Rahnenfuhrer J (2020)<sup>48</sup> | 2.4.0 |
| biomaRt | Durinck et al (2009)<sup>49</sup> | 2.40.5 |
| scRNA-seq for Mouse Nervous System | Zeisel et al. (2018)<sup>50</sup> | http://mousebrain.org/ |
| Ligand-Receptor pair database | Ramiloski et al (2015)<sup>51</sup> | https://fantom.gsc.riken.jp/5/suppl/Ramilowski_et_al_2015/ |

#### Single cell RNA-sequencing using 10X Genomics platform

Single-cell suspensions were prepared as described above. Two biological replicates for each time point for a total of 8 samples were sequenced with a median of ~8000 cells per sample. The median of mean reads per cell across samples was > 55,000 and median sequencing saturation was 81.2%. Libraries for all samples were prepared according to Chromium Single Cell 3’ Library and Gel Bead Kit v2 instructions (10X Genomics, PN-120237) and indexed with Chromium i7 Multiplex Kits (10X Genomics, PN-120262). In brief, each cell suspension was first partitioned into single-cell gel beads-in-emulsion (GEMs) and then incubated with reverse-transcriptase primers and reaction mix. Primers contain i) a 16nt 10X barcode, ii) a 10nt Unique Molecular Identifier, iii) a poly-dT primer sequence, and iv) an Illumina Read 1 (R1) sequence to produce single-stranded, barcoded complementary DNA (cDNA) from poly-adenylated mRNA. After breaking GEMs to recover cDNA fractions, leftover biochemical reagent was removed with Silane Dynabeads (10X Genomics, PN-2000048), and pooled fractions were amplified via PCR to generate full-length, double-stranded, barcoded cDNA. Products were then enzymatically fragmented, end repaired, and modified with dA-tails. Insert sizes were filtered with double-sided size selection using SPRIselect (Beckman Coulter, B23318) before cDNA molecules were ligated to Illumina Read 2 (R2) adapters. To construct final libraries for Illumina sequencing, P5, P7, and sample index sequences were inserted into molecules by sample index PCR and then amplified. For sequencing, libraries were loaded at recommended loading concentrations onto an Illumina NextSeq 500 flow cell and paired-end sequenced under recommended settings (R1: 28 cycles; i7 index: 8 cycles; i5 index: none; R2: 91 cycles). After sequencing, Illumina output was processed using CellRanger’s recommended pipeline to generate gene-barcode count matrices. In brief, base call files for each sample were demultiplexed into FASTQ reads and then aligned to the mouse mm10 reference genome using STAR splice-aware aligner. Reads that confidently intersected at least 50% with an exon were considered exonic and further aligned with annotated transcripts. Reads were then filtered to remove UMIs and barcodes with single base substitution errors and finally used for UMI counting. The output was a count matrix containing all UMI counts for every droplet. CellRanger v2.2.0 was used for the first set of replicates and v3.0.1 for the second set of replicates.

#### Preprocessing and Quality Control

To distinguish cell-containing droplets from empty droplets, we performed cell calling on the unfiltered UMI count matrices using a combination of barcode-ranking and empty-droplet detection algorithms. First, cell barcodes were ranked according to UMI count and visualized in a log-total UMI count vs log-rank plot. A spline curve was fit to the data to identify knee and inflection points. At least all data points above the knee were considered cell-containing droplets. In order to further distinguish cells from data below the knee, we used the emptyDrops function from the DropletUtils R package^52^ using the following fixed parameters for all sample: lower = 250; max fit.bounds = 1e06; FDR = 0.001; ignore = 10. Some parameters varied between samples accordingly: “retain” was set to knee point values, and “lower” was set to inflection point values. The result was a filtered count matrix of all putative cell-containing droplets. In order to distinguish low quality cells, we considered cell-level metrics such as library size, percentage of UMIs mapping to mitochondrial genes, and doublet detection algorithm outputs. We observed that removing cells in the lowest quantiles of total UMI counts preferentially selected against endothelial cells, while removing cells in the highest quantiles of mitochondrial percentage preferentially selected against astrocytes. To avoid bias due to any one metric, we used a multivariate approach to automatic outlier detection as implemented in the Scater R package^45^, which performs principal component analysis on the quality control metrics to determine outliers.

In order to remove potential doublets, we applied the Scrublet Python package^47^ for each individual sample. In brief, the Scrublet simulates multiplets by sampling from the data and builds a nearest-neighbor classifier. Cells from the data that have high local densities of simulated doublets are flagged and removed. We set the expected_doublet_rate for each sample according to the estimated doublet rate per cells sequenced as published by 10X, and default values for all other parameters. During downstream analysis, we observed small, unidentifiable clusters that co-expressed marker genes for two different cell-types, and removed these cells as they were likely to be multiplets not detected by Scrublet. Additionally, we identified a cluster enriched for hemoglobin genes (Hbb-bs, Hbb-bt, etc.) which was removed from downstream analyses.

#### Normalization and Batch Correction

To generate the full SCI dataset, all samples were processed and combined using Seurat v3^53^. After filtering each sample count matrix for genes that were expressed in at least 10 cells, each dataset was independently normalized and scaled using the SCTransform function. SCTransform performs a variance-stabilizing transformation in which genes are grouped according to mean expression in order to smoothen model parameters for negative binomial regression^54^. To remove cell-cycle genes as a confounding source of variation, mitochondrial percentage and cell cycle scores based on the expression of canonical G2M and S phase markers were computed for each cell. Cell cycle genes were provided through the Seurat tutorial. These score values were then used as input for the “vars.to.regress” argument in the SCTransfrom() function. This operation generates a “corrected” expression matrix by building a regression model on these variables for each gene. To identify shared and unique molecular cell-types across datasets and time-points, sample expression matrices were batch-corrected using Seurat’s Data Integration workflow^55^. In brief, data integration uses canonical correlation analysis between two samples to identify mutual nearest neighbors. These “anchors” are then used to generate a corrected gene expression matrix based on the consistency of the anchors between cells, effectively performing a batch correction. For the full SCI dataset, the 2000 most variables genes were used as input for the “anchor. features” argument of the FindIntegrationAnchors() function, where the variance of a gene was measured as the residual divided by the expected variance under the SCTransfrom() model. This resulted in a single, batch-corrected expression matrix for containing all cells.

For the analysis of the myeloid cells, we tested a similar batch correction as described above. However, we observed that some microglia from the uninjured spinal cord were classified under a peripheral macrophage subset but not under BA-Macrophage (data not shown). We attributed this potential misclassification to the imbalance between uninjured and injured cell counts, and the large proportion of peripheral myeloid cells in the injured samples. We instead combined sample datasets across time-points for each replicate and subsequently performed normalization for each of the two larger datasets as above (i.e. normalize across times and batch-correct across replicates). For myeloid integration, the 2000 most variable genes were used. For the vascular cells, no obvious misclassifications were noted. We proceeded with normalization and batch correction as above, using the 2000 most variable genes for downstream analysis. For the macroglia cells, we again observe no obvious misclassifications and proceeded with normalization and batch correction. However because of the low numbers of macroglia cells in one of the uninjured sample replicates, we adjusted the “k.filter” parameter to 100 for the FindIntegrationAnchors() function.

#### Dimensional Reduction, Clustering, and Differential Gene expression testing

In order to measure the greatest gene expression variation among all the SCI cells, we first performed PCA on the batch-corrected expression matrix for the top 2000 variable genes taken from above. The top 15 principal components (PCs) were selected based on the “elbow” point heuristic in a scree plot which quantifies the contribution of variance by each principal component. Using these components, a nearest-neighbor graph and shared-nearest-neighbor graph were generated with “k.neighbors” set to 20 by default. To visualize the cells, we generate a UMAP plot with default Seurat parameters using cell coordinates in PCA-space using the top 15 PCs. In order to cluster the cells based on similarity of expression, we ran the FindClusters() function on the shared-nearest-neighbor graph with default parameters.

For the myeloid, vascular, and macroglial cells, we performed similar analyses as described above, with a few modifications. In order to identify reproducible sub-clusters of cells, we performed the same graph-based clustering through a range of PCs, “k.neighbor” and “resolution” parameters and inspected cluster memberships for stable configurations. For the myeloid, vascular, and macroglia, we took the top 12, 11, and 8 PCs and set resolutions to 0.5, 0.3, and 0.45 respectively. To identify marker genes for each cluster, we used the FindAllMarkers() function using default parameters, which implements a Wilcoxon Rank Sum test comparing gene expression of cells within a given cluster versus all other cells. We repeated this analysis to identify marker genes distinguishing subsets within a cell-type.

To infer the functional relevance of sub-clusters, we performed gene ontology enrichment analyses on the top 50 differentially expressed genes using Fisher’s Exact test as implemented in the topGO R package^48^. For the enrichment analyses of the gene expression changes in astrocytes, our initial analysis revealed very few differentially expressed genes between the uninjured and 1dpi astrocytes, which we attributed to the low numbers of uninjured astrocytes captured. Therefore, we supplemented our uninjured astrocyte dataset with ACNT1 and ACNT2 astrocyte data from the previously published mouse CNS single-cell atlas dataset^50^. We also supplemented our uninjured OPC dataset in order to validate that our uninjured cells were more transcriptional similar to the external reference cells than to our injured cells. To account for differences in sequencing depth between our dataset and the external dataset, we performed differential expression tests using MAST as implemented in Seurat^56^. We used all differentially expressed genes (p_val_adj < 0.001) as input for gene ontology analysis.

#### Ligand-Receptor Interaction Analysis

To infer potential ligand-receptor interactions between two cell-types, we adapted the method used in CellPhoneDB^15^. We first pulled a reference list of human ligand-receptor pairs published previously^51^ and converted the genes into mouse orthologs using the Ensembl biomaRt package^49^. We defined the ligand-receptor score as the mean of the average log-normalized expression of the receptor gene in one cell-type and the average log-normalized expression of the ligand gene in a second cell-type. To identify enriched ligand-receptor interactions, we applied a permutation test to identify interactions scores that are enriched in a specific <ligand cell A, receptor cell B, time-point> combination. For each of 1000 permutations, we randomly shuffled the cell-type and time-point labels and calculated an interaction scores for all possible <ligand cell, receptor cell, time-point> combinations. Repeating this 1000 times generated a null distribution of interaction scores for each ligand-receptor pair. We compared the interaction scores of the actual (ligand cell A, receptor cell B, time-point) labels to the null distribution and calculated p-values as the proportion of null scores which are equal to or greater than the actual interaction score.

## Notes

### Competing Interest Statement

The authors have declared no competing interest.

