## Supplemental figures for "Single cell analysis of the cellular heterogeneity and interactions in the injured mouse spinal cord"

**a**

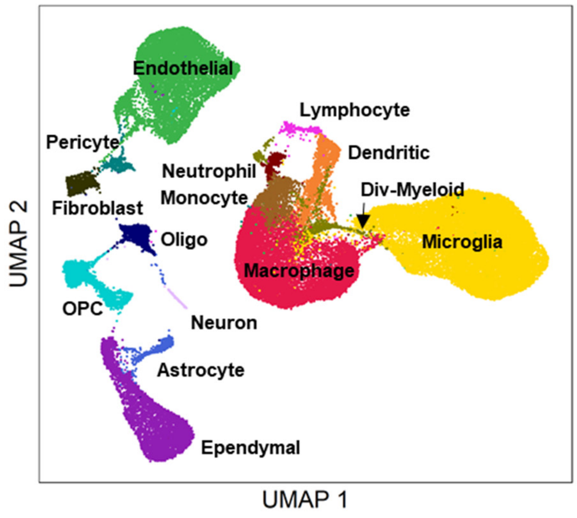


**b**

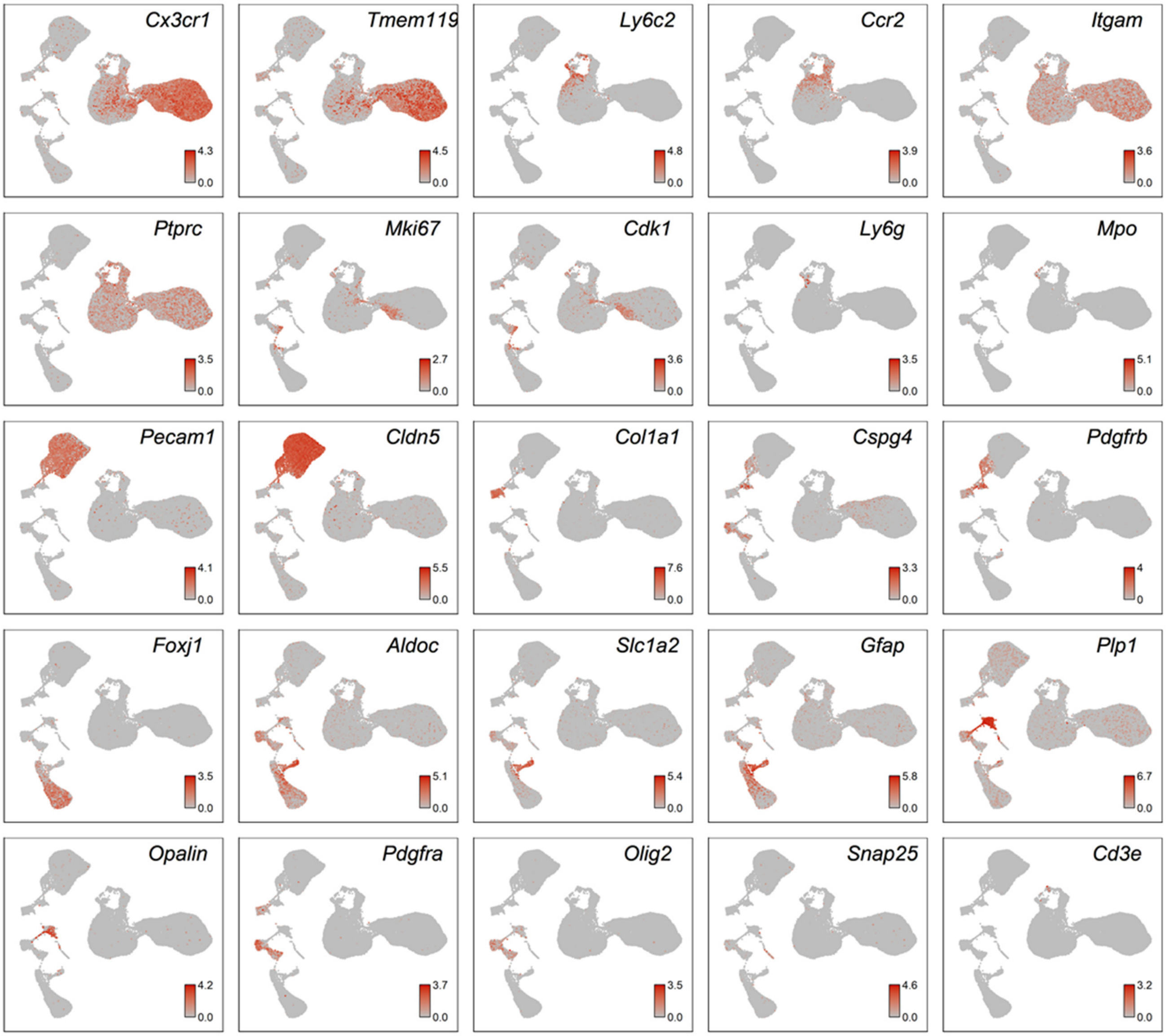


Extended Data Fig. 1

**Extended Data Figure 1| Expression pattern of canonical genes used to identify major cell types.** (a) UMAP visualization of all cells from all time points. (b) Expression pattern of previously annotated marker genes used to identify major cell types on the UMAP. Cx3cr1 and Tmem119 identified microglia. Itgam and Ptprc identified leukocytes. Ly6C2 and Ccr1 identified monocytes. Ly6G identified neutrophils. Cd3e identified lymphocytes. Mki67 and Cdk1 identified dividing cells. Pecam1 and Cldn5 identified endothelial cells. Col1a1 identified fibroblasts. Combined expression of Cspg4 and Pdgfrb identified pericytes. Foxj1 identified ependymal cells. Combined expression of Aqp4 and Gfap identified astrocytes. Plp1 and Opalin identified oligodendrocytes. Combined expression of Pdgfra and Olig2 identified oligodendrocyte progenitor cells. Snap25 identified neurons.

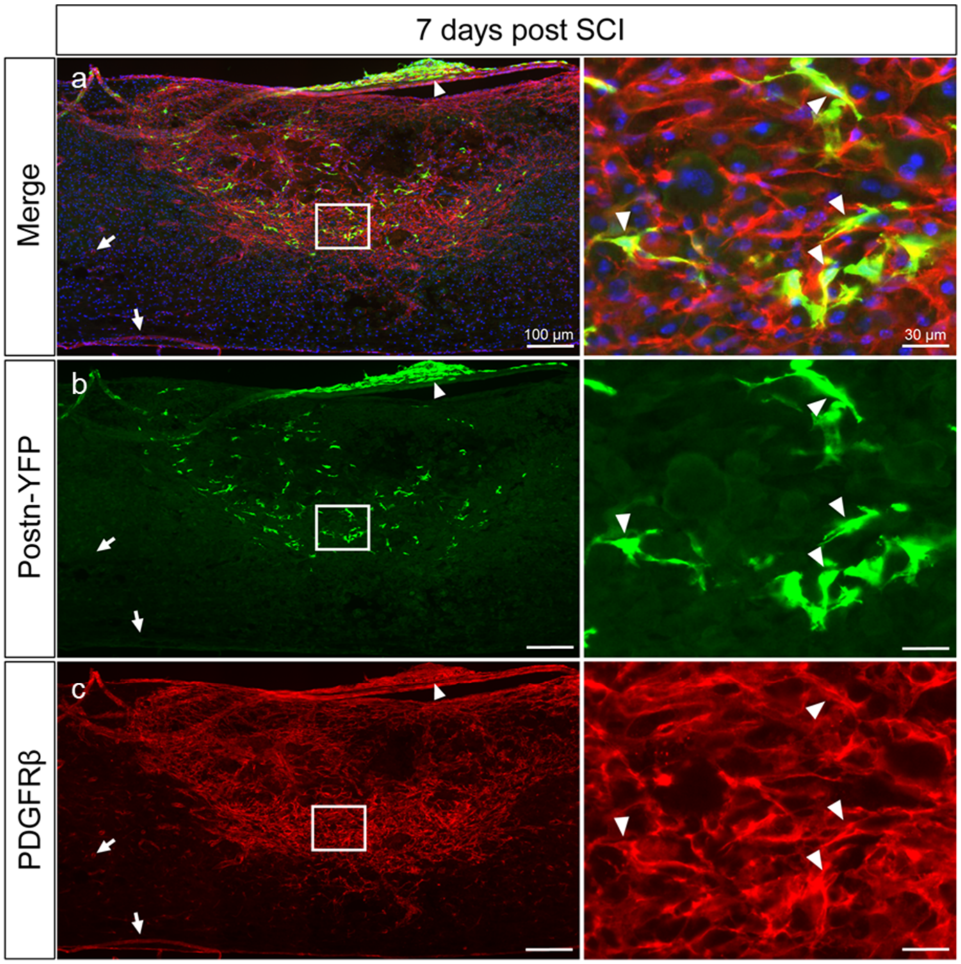


Extended Data Fig. 2

**Extended Data Figure 2| Validation of periostin as a marker of activated fibroblasts after SCI.** Immunohistochemical analysis of 7dpi spinal cord from Postn-CreER/Rosa26-EYFP mice show genetically labeled EYFP+ fibroblasts (b) exclusively located in the fibrotic area delineated by dense PDGFR-β+ region as well as only in meningeal fibroblasts that overly the injury site (c). Arrow heads denote activated fibroblasts that are both EYFP+/ PDGFR-β+. PDGFR-β+ fibroblasts in the fibrotic scar that do not express EYFP are due to low recombination efficiency in these mice. Arrows indicate perivascular PDGFR-β+ cells in uninjured regions that do not express EYFP. Boxed regions are magnified on the right panels. Scale bar on left = 100µm. Scale bar on right = 30µm

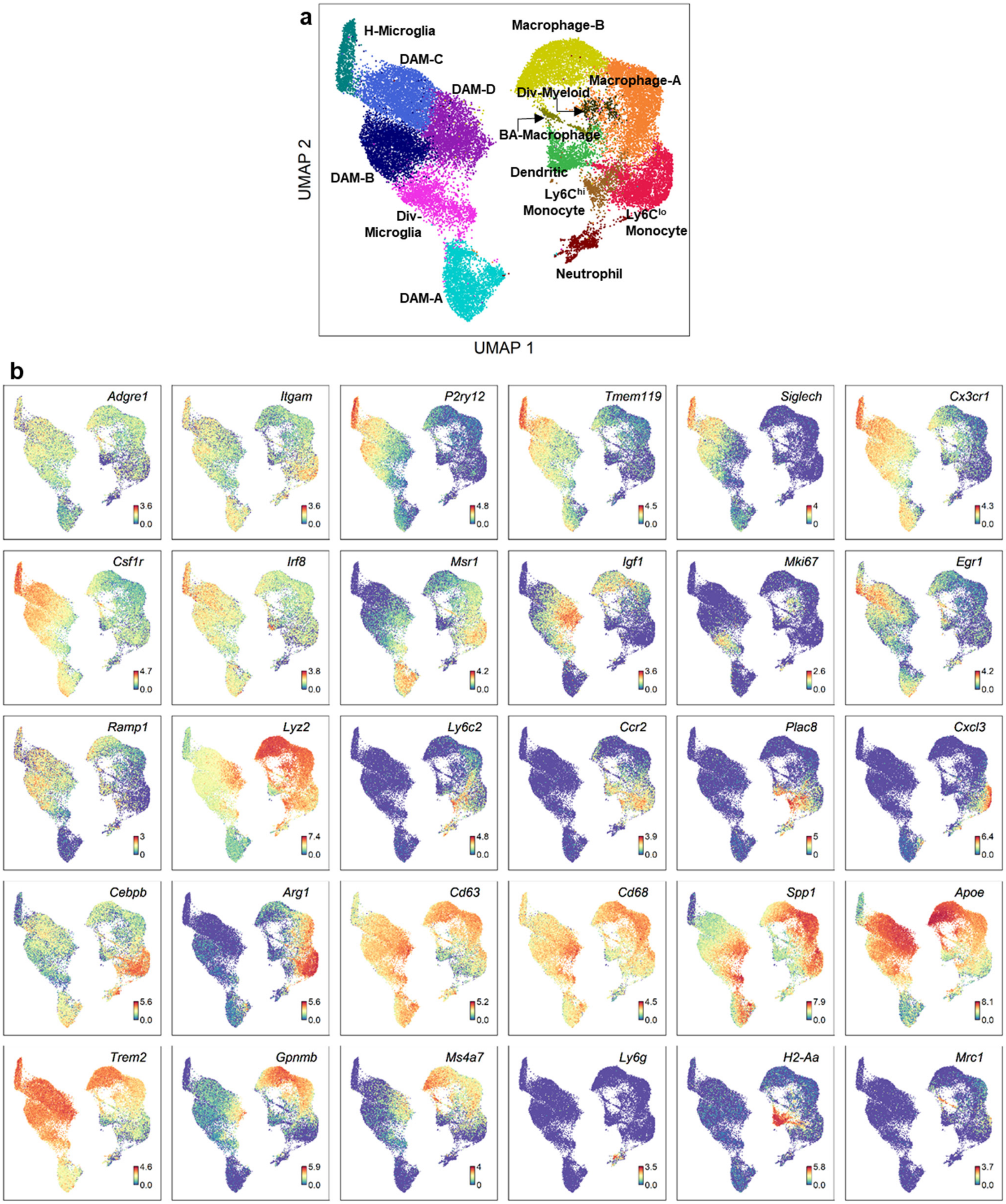


Extended Data Fig. 3

**Extended Data Figure 3| Expression pattern of annotated genes used to identify myeloid cell types.** (a) UMAP visualization of all myeloid cells from all time points. (b) Expression pattern of previously annotated marker genes used to identify myeloid cell types on the UMAP. The large cluster on the left was identified as microglia by the expression of *p2ry12*, *tmem119*, and *siglech*. The large cluster on the right was identified as peripheral myeloids by the expression of *ccr2* and *ly6c2*. Homeostatic microglia (H-microglia) was identified by high expression of *p2ry12*, *siglech*, and *tmem119*. Disease-associated microglia (DAM) were identified by expression of *lpl*, *msr1*, and reduced expression of the homeostatic microglia markers. Dividing myeloid cells were identified by *mki67* and *top2a*. Monocytes were identified by *ccr2* and *ly6c2*. Macrophages were identified by *cd63* and the reduced expression of monocyte markers.

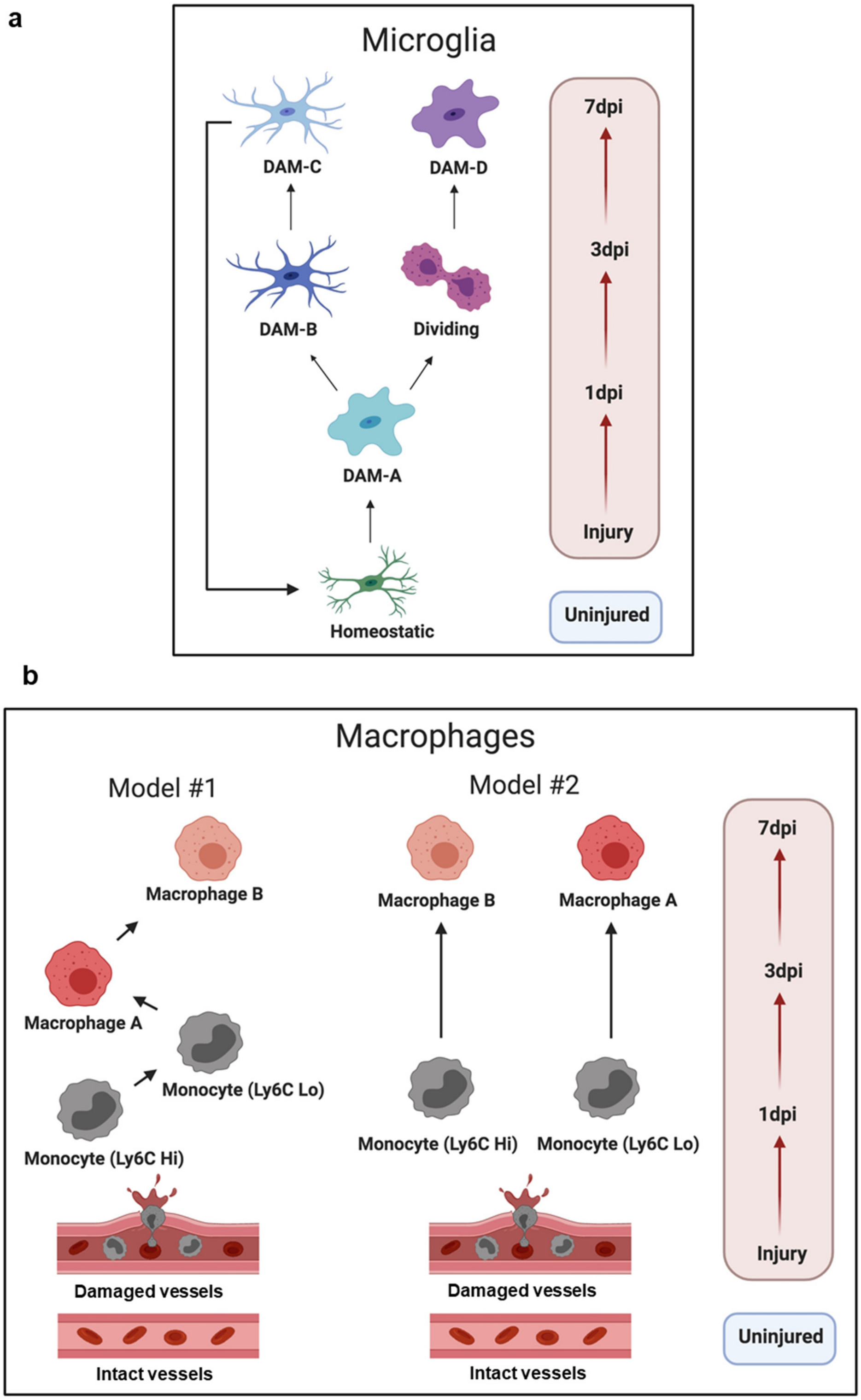


Extended Data Fig. 4

**Extended Data Figure 4| Models of microglia and macrophage state dynamics after SCI.** (a) After injury, homeostatic microglia become Disease-Associated Microglia subtype A (DAM-A). Some DAM-A proliferate and give rise to DAM-D, while others return to a more homeostatic state through DAM-B and DAM-C. (b) Two models of monocyte-macrophage differentiation are possible. In the linear model (left), Ly6C$^{hi}$ monocytes enter from the blood and become Ly6C$^{lo}$ monocytes, which differentiate into Macrophage-A and then to Macrophage-B that persists chronically in the injured spinal cord. In the parallel model (right), Ly6C$^{hi}$ and Ly6C$^{lo}$ monocytes enter as distinct populations and each give rise to Macrophage-B or Macrophage-A.

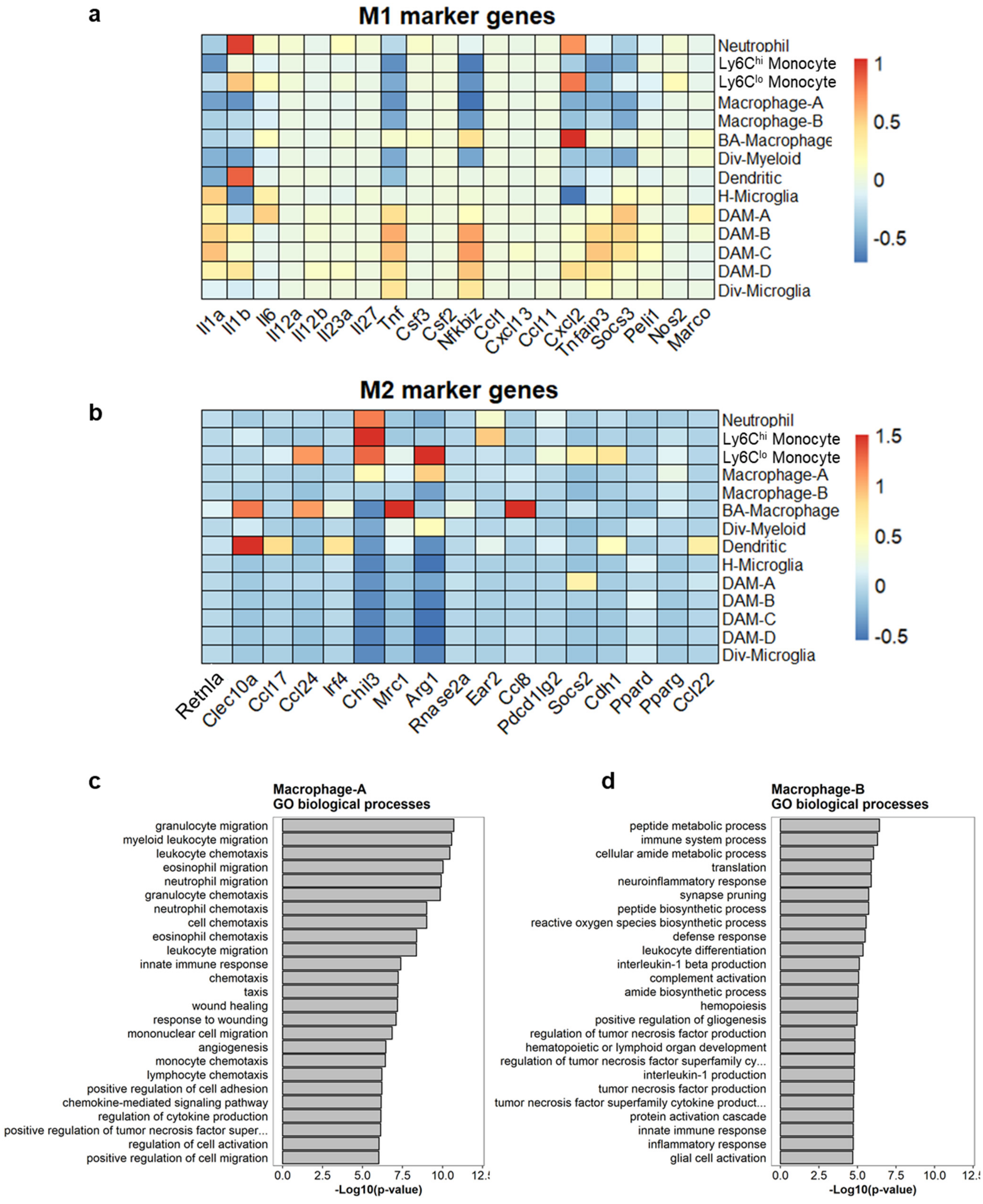


Extended Data Fig. 5

**Extended Data Figure 5| Comparison of Macrophage-A and B subtypes based on M1/M2 markers and Gene Ontology (GO) terms.** Heat map of annotated M1 (a) and M2 (b) macrophage gene expression shows that neither Macrophage-A nor Macrophage-B subtypes display distinct M1/M2 nomenclature compared to other myeloid cells. GO biological processes based on highest differentially expressed genes in Macrophages-A (c) and Macrophage-B (d).

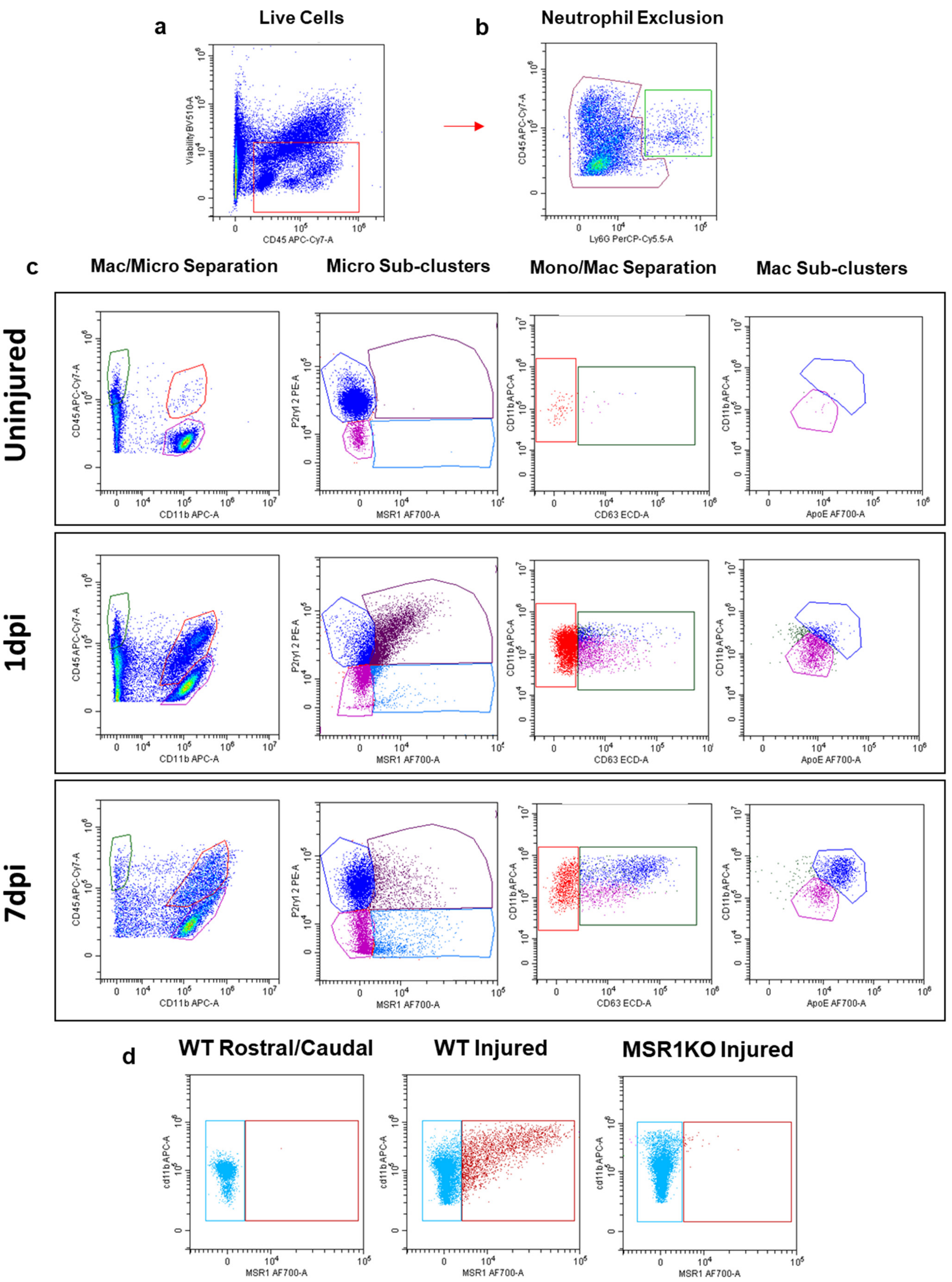


Extended Data Fig. 6

**Extended Data Figure 6| Gating strategy for flow cytometry validation of myeloid cell Subtypes.** After selecting for viable $CD45^+$ leukocytes (a), neutrophils were excluded based on Ly6G expression (b). (c) $Ly6G^-$ cells were separated based on CD11b and CD45, which identified lymphocytes ($CD11b^-$), monocyte/macrophages ($CD11b^+/CD45^{hi}$), and microglia ($CD11b^+/CD45^{lo}$). The microglia cluster was gated on P2ry12 and Msr1 to identify DAM-A. The monocyte/macrophage cluster was first gated on $CD63^+$ cells to identify macrophages, and then gated on ApoE and CD11b to separate Macrophage-A and B subtypes. (d) Spinal cord tissue rostral and caudal to the injury site as well as injury site from Msr1 knockout mice were used as negative controls for Msr1 gating.

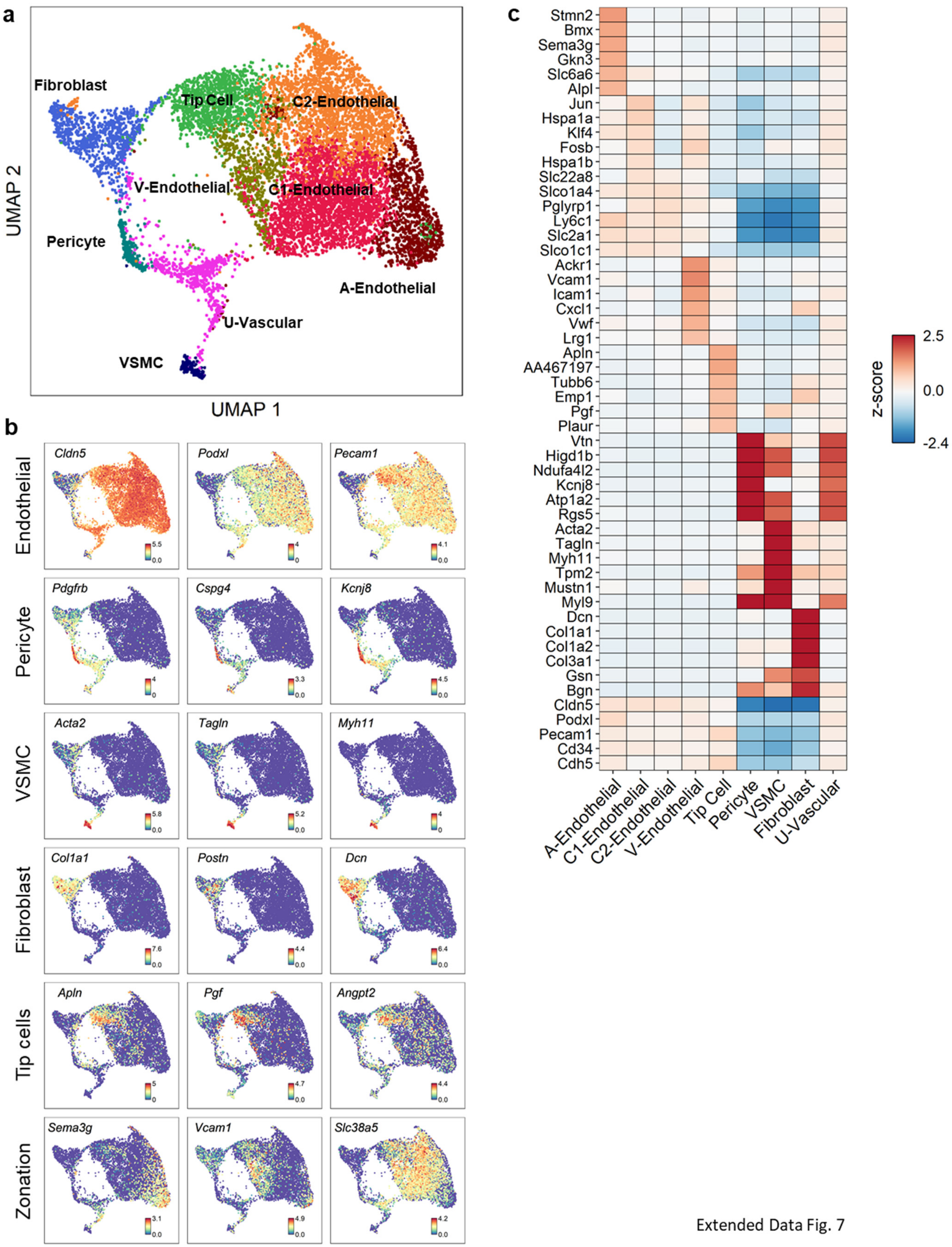


Extended Data Fig. 7

**Extended Data Figure 7| Molecular profile of vascular cells after SCI.** (a) UMAP clusters of all vascular cells from all time points. (b) Expression pattern of previously annotated marker genes used to identify each cluster. (c) Heatmap of differentially expressed genes for each cell type.

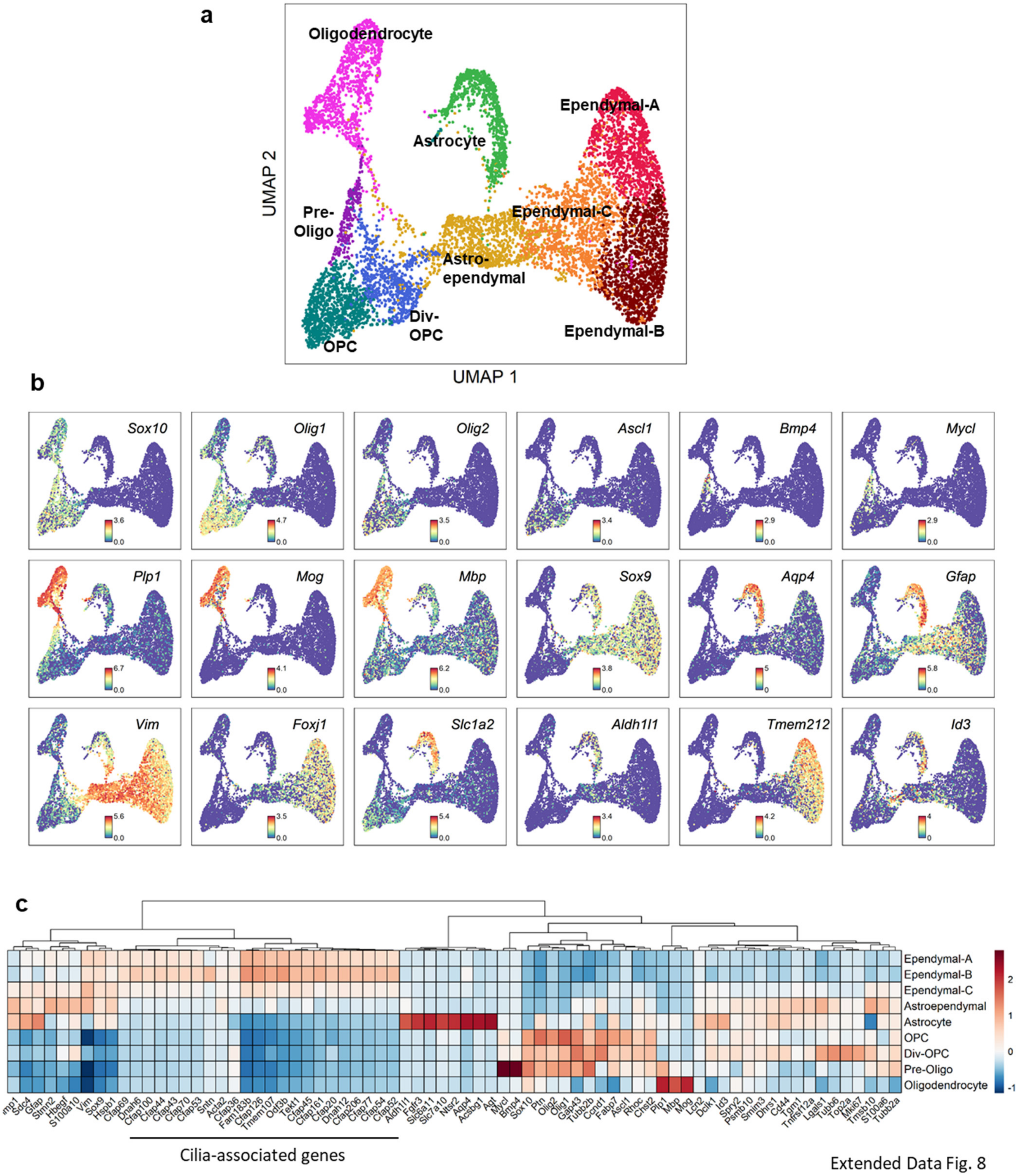


Extended Data Fig. 8

**Extended Data Figure 8| Molecular profile of macroglia cells after SCI.** (a) UMAP clusters of all macroglia cells from all time points. (b) Expression pattern of previously annotated marker genes used to identify each cluster. (c) Heatmap of astrocyte and ependymal markers to assess their similarities and differences to astroependymal cells.

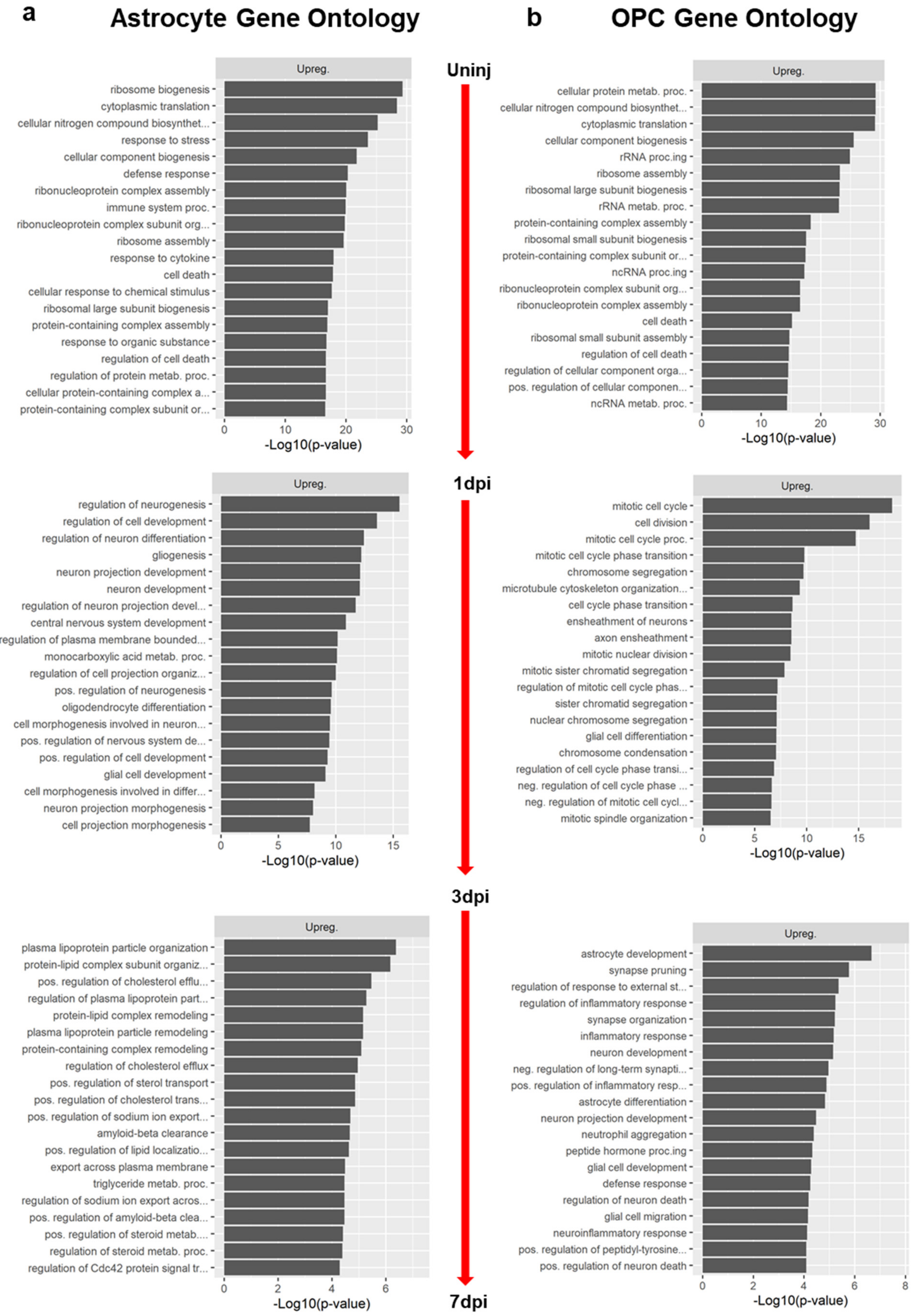


Extended Data Fig. 9

**Extended Data Figure 9| Comparison of Gene Ontology (GO) biological processes between astrocytes and oligodendrocyte progenitors cells (OPCs).** GO biological processed based on the highest DEGs between sequential time points between astrocytes (a) and OPCs (b) after SCI. The top five terms at each time point are shown in Fig. 6. The remaining terms are shown here.
